# Cognitive exertion reshapes resting-state EEG markers across the adult lifespan

**DOI:** 10.64898/2026.08.14.744886

**Authors:** Kitti Bán, Béla Weiss, Tibor Auer, Patrick D. Gajewski, Edmund Wascher, Zoltán Vidnyánszky

**Affiliations:** Brain Imaging Centre, HUN-REN Research Centre for Natural Sciences, Budapest, Hungary; Machine Perception Research Laboratory, HUN-REN Institute for Computer Science and Control, Budapest, Hungary; Faculty of Brain Sciences, Information Services Division, University College London, London, United Kingdom; Department of Ergonomics, Leibniz Research Centre for Working Environment and Human Factors at TU Dortmund, Germany; German Center for Mental Health (DZPG), partner site Bochum/Marburg, Germany

**Keywords:** EEG, ageing, individual alpha oscillations, aperiodic exponent, cognitive exertion, trait, state

## Abstract

Resting-state electroencephalography (rsEEG) yields robust indices of ageing, notably individual alpha peak frequency (iAPF), alpha power, and the aperiodic exponent. Whether these markers reflect stable traits or shift with cognitive exertion remains unresolved, with direct consequences for lifespan and clinical research. We parameterised periodic and aperiodic rsEEG activity before and after cognitive tasks in a lifespan cohort (*N* = 390, aged 20–70), a five-year longitudinal follow-up (*N* = 100), and an independent older-adult dataset completing a different task (*N* = 71). Across datasets, cognitive exertion produced lifespan-wide iAPF slowing and increase in associated power. Notably, aperiodic shifts were age-dependent, with post-task steepening of the exponent in younger adults that progressively flattened with advancing age, resulting in a stronger effect of age on post-task exponents. Our results demonstrate that widely used spectral metrics exhibit acute state-dependency and post-task recordings offer a promising translational framework for indexing individual differences in healthy ageing and pathology.

**Trial registration:** Clinicaltrials.gov: NCT05155397

## Introduction

Mapping the pervasive functional reorganisation of the ageing brain non-invasively across the lifespan is a central pursuit of cognitive and translational neuroscience. The brain’s spontaneous electrical activity during wakeful rest provides a valuable window into these processes as both normative ageing and age-related pathologies manifest as altered dynamics within large-scale neural networks detectable via resting-state recordings^1^. Compared to functional magnetic resonance imaging (fMRI), resting-state electroencephalography (rsEEG) offers uniquely scalable, mobile, and cost-effective insights into these neurophysiological shifts^2^. As a result, a variety of rsEEG-derived metrics have become adopted indices of individual differences in brain ageing and neuropathology^3,4^.

Spectral features of rsEEG have proven especially valuable in this context, owing to their computational simplicity^5^ and excellent reproducibility^6,7^. Historically, such work characterised oscillatory activity within canonical frequency bands, including delta (1–4 Hz), theta (4–8 Hz), alpha (8–12 Hz), beta (13–30 Hz), and gamma (30–60 Hz). These frequency band measures, typically peak frequency and power, have been repeatedly linked to cognition^8,9^, developmental and ageing trajectories^10–14^ and neuropathology, including depression^3,15^, anxiety^16^, and Alzheimer’s disease^4,17^. Among spectral features, alpha band metrics, primarily individual alpha peak frequency (iAPF) and related power, consistently index lifespan shifts^11–13,18^ positioning them as standard metrics in neurophysiological models of ageing.

Within the EEG power spectrum, oscillatory activity is superimposed on a background aperiodic component, which exhibits an 1/f-like power decrease with increasing frequency^19,20^ (Fig.1). This aperiodic component consists of the aperiodic exponent, which indexes the slope of spectral decay in log–log space, and the aperiodic offset, which represents broadband power across the spectrum. Critically for ageing research, the exponent is commonly interpreted as a proxy for cortical excitation and inhibition (E/I) balance^19,21,22^, and its flattening across the adult lifespan has been proposed to reflect an age-related shift towards relative excitation^18,20,22^. Moreover, exponent alterations have also been reported in neuropathology, including dementia^23^, depression, and ADHD^24^, establishing it as a promising marker of both healthy ageing and neuropathology. Given the substantial shared variance between the two aperiodic parameters, the offset frequently conveys information parallel to that of the exponent^18,25,26^.

**Figure 1.**
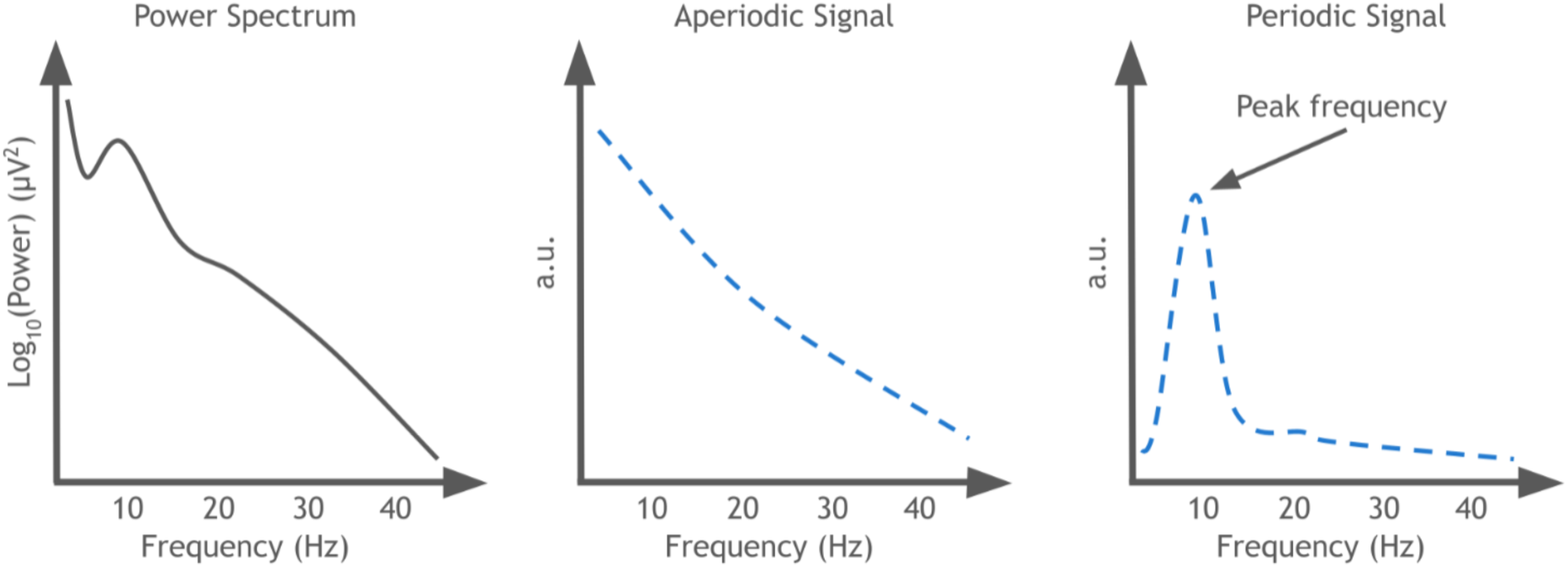
Schematic representation of spectral parameterisation. The overall EEG power spectrum (left) is decomposed into its underlying aperiodic (exponent and offset; middle) and oscillatory, periodic (right) components.

A core premise underlying the application of periodic and aperiodic markers in ageing and clinical research is that these features represent stable, trait-like neurophysiological characteristics^27^. However, emerging evidence indicates that resting-state spectral measures are sensitive to state-dependent modulation following acute cognitive exertion. In young adults, demanding cognitive tasks have been reported to induce an increase in resting-state power^28–30^ alongside a concurrent decrease in iAPF^28,31^. Crucially, the few available studies in older populations report divergent outcomes, describing post-task power increases^30^, decreases^32^, as well as null effects^33^. These differences raise the possibility that the ageing brain’s spectral response to cognitive demand differs systematically from that of younger adults.

However, evidence to support this conjecture is sparse and inconsistent, with the majority of existing investigations failing to separate periodic from aperiodic activity when quantifying task-related changes. It is therefore uncertain whether reported power alterations stem from genuine oscillatory shifts, an aperiodic redistribution, or a combination of the two. To our knowledge, Yang et al.^30^ represent the sole study to have decomposed the spectrum and evaluated task-induced shifts in both younger and older cohorts. Specifically, they observed a post-task increase in the aperiodic exponent in both groups, an effect that was attenuated in the older cohort. Nonetheless, this study relied on eyes-closed recordings and utilised categorical age splits, precluding the examination of continuous lifespan trajectories underlying task-related spectral shifts.

Resolving these questions matters for how rsEEG is utilised in ageing research. Traditionally, for spectral markers to reliably index trait-like variables such as chronological age or neurophysiological decline, their measurement properties are assumed to require robustness against transient, within-session fluctuations^27^. An alternative perspective suggests that rather than introducing mere measurement noise, task-induced state transitions may act as a neurophysiological stress-test, unmasking underlying individual differences that remain latent during baseline conditions^34,35^. This latter view aligns with evidence indicating that compared to pre-task baselines, post-task EEG metrics are superior at differentiating between clinical and healthy cohorts, including individuals with brain injury or cognitive impairment^32,34,36,37^. Extending this framework to a lifespan perspective, if the degree of such state-dependent shifts varies continuously with age, the relationship between spectral metrics and age will systematically differ depending on whether they are sampled at baseline or following cognitive exertion. Consequently, we hypothesised that age would exhibit a stronger effect on post-task metrics compared to their baseline, pre-task equivalents.

Addressing this hypothesis has recently been made possible by the Dortmund Vital Study^38^ (DVS), which overcomes previous dataset limitations by providing pre- and post-task rsEEG across over 600 healthy adults aged 20–70. Crucially, this cohort combines eyes-open and eyes-closed conditions as well as a 5-year longitudinal follow-up for approximately half of the sample. Leveraging these data, the present study characterises how acute cognitive exertion reshapes prominent rsEEG spectral metrics across the adult lifespan, whilst evaluating the temporal stability and cross-paradigm generalisability of these shifts.

## Methods

### Participants

#### Dortmund Vital Study

We utilised a large, longitudinal resting-state EEG dataset collected as part of the longitudinal Dortmund Vital Study^38,39^ (for more details, see the study protocol registered as NCT05155397 on <u>ClinicalTrials.gov</u> or the protocol paper^39^). Session 1 data include recordings from 623 participants (age range 20–70 years, mean (*M)* = 43.79 years, standard deviation (*SD)* = 14.45 years; 388 female). All participants were invited for a follow-up session (Session 2) approximately 5-years following the initial assessment, of whom 329 participants (Session 1 age: *M* = 43.79 years, *SD* = 14.45 years, 188 female) participated during this second time point.

Participants exclusion criteria included a history of major diseases (for further details, see Getzmann et al.^38^), the use of psychotropic drugs and narcoleptics, as well as limited physical fitness and mobility. The study was approved by the Ethical Committee of the Leibniz Research Centre for Working Environment and Human Factors, Dortmund, Germany (approval number: A93-1 and A93-3 for the follow-up testing) and conformed to the Declaration of Helsinki. All participants provided written, informed consent prior to participation. Demonstrating the technical and psychometric rigorousness of these data, spectral metrics within the DVS have been reported to show excellent long-term test-retest reliability^6,7^.

Following data pre-processing and applying related exclusion criteria (see ‘EEG pre-processing’ and ‘Calculation of periodic and aperiodic metrics’), we utilised a stratified random sampling approach in order to ensure a balanced distribution of age and gender across the Session 1 and 2 datasets. First, participants were categorised into five, mutually exclusive 10-year age bins. Within each bin, the data were further stratified by sex (male and female). To achieve bin equality, the subgroup with the lowest frequency across all age-by-gender combinations was identified as the baseline constraint. Finally, a random selection process was executed within each stratum to match this minimum threshold, resulting in an identical number of male and female participants across all age intervals. For Session 1, 39 males and females were included in each age bin, resulting in a total of 390 participants (*M* = 44.59, *SD* = 14.33 years). For Session 2, 10 males and females were selected for each stratum, resulting in a total of 100 participants (M = 49.80, SD = 14.90 years). Our longitudinal analyses contained data from the same 100 participants that were selected for the Session 2 analyses.

#### Validation dataset

Data from 78 people between the ages 63-77 years (*M* = 70.02, *SD* = 3.22, 58 females) were obtained at the Brain Imaging Centre within the HUN-REN Research Centre for Natural Sciences. All participants had corrected-to-normal vision. Exclusion criteria included pre-existing major medical disorders (e.g., cognitive impairment, neurological or psychiatric conditions, cardiovascular disease, etc.) and medication use affecting cognition or movement. The study was conducted in accordance with the principles of the Declaration of Helsinki and the protocol was approved (approval number: NNGYK/50614-5/2024) by the National Centre for Public Health and Pharmacy (Budapest, Hungary). All participants provided written, informed consent prior to participation.

### Resting-state EEG acquisition

#### Dortmund Vital Study

For all participants, data collection during both sessions took part during the same time in the morning to promote a state of wakefulness. Eyes-open and eyes-closed resting-state data were collected for 3 minutes each before and after an approximately 2-hour cognitive task battery. Eyes-closed recording always took place prior to the eyes-open recordings. The test battery consisted of a total of five tasks assessing basic cognitive functions, comprising (1) selective attention and perceptual control, (2) sustained attention (Psychomotor Vigilance Test), (3) conflict processing (Simon Task), (4) cognitive control and context updating (AX-Continuous Performance Task), and (5) speech perception under auditory distraction (Speech-in-Noise Perception Task). The tasks had a medium to high level of difficulty (for details, see Getzmann et al.^38^ and Gajewski et al.^39^). The exact same procedure was followed for Sessions 1 and 2.

Scalp EEG was recorded using a 64-channel actiCAP active electrode system (BrainProducts GmbH, Germany) positioned following the international 10-20 system, with electrode FCz as an on-line reference. Data were amplified using a BrainAmp DC amplifier (BrainProducts GmbH, Germany) and recorded in Brain Vision Recorder (Brain Products, Germany) at 1000 Hz and filtered online by a 250-Hz low-pass filter. Electrode input impedance was adjusted to under 10 kΩ.

#### Validation dataset

Upon arrival, participants provided informed consent and were asked about the number of hours slept, their sleep quality over the previous night, and whether they felt overly satiated from a recent meal. In cases of little/poor sleep or over-satiation, the experimental session would be rearranged to another day, however, there were no such instances.

Simultaneous resting-state EEG and eye-tracking data were recorded for 3 minutes before and after an approximately 1.5-hour-long probabilistic reversal learning task^40^. During the resting-state period, participants were instructed to fixate on a black cross in the middle of a grey screen, whilst remaining still and sitting comfortably with their chin positioned in a height-adjustable chin rest 75 cm from the display (24’’ Fujitsu LED monitor, 60 Hz refresh rate) in a dark, soundproof, and electrostatically shielded room. No specific instructions on blinking were provided.

Scalp EEG was recorded using a 64-channel actiCAP active electrode system (BrainProducts GmbH, Germany) situated following the international 10-10 system and amplified using a BrainAmp DC amplifier (BrainProducts GmbH, Germany). The ground and reference electrodes were located at AFz and FCz, respectively. Electrode input impedance was adjusted to under 10 kΩ. Data were recorded in Brain Vision Recorder (Version 1.20, Brain Products, Germany) at 1000 Hz and subjected to online (hardware) filtering by an digital band-pass filter of 0.01 - 250 Hz. Experimental trigger codes marking the start and end of the 3-minute resting period were synchronised between the EEG and eye-tracking data. We utilised eye-tracking to validate participant wakefulness during resting-state recordings (detailed acquisition parameters are described in the Supplementary Material).

### EEG pre-processing

We utilised an identical pre-processing approach for all datasets. All EEG datasets were organised and converted to the Brain Imaging Data Structure^41^ (BIDS) format. Raw EEG data pre-processing was conducted in Automatic Analysis^42^ (version 5.8.1), EEGLab^43^ (version 2024.2), and FieldTrip^44^ (git revision 2755b10) running on MATLAB (version 2024b, The Mathworks Inc., 2024).

First, data were down-sampled to 250 Hz. We implemented a high-pass filter at 1 Hz, a low-pass filter at 120 Hz, as well as band-stop filters at 45-55 Hz and 95-105 Hz. To remove artefactual channels and data points, we utilised the clean_rawdata plugin within EEGLab^45^ (version 2.10). We rejected channels with a flatline of longer than 5 seconds or when correlation with nearby channels was below the 0.8 cut-off value. We used Artifact Subspace Reconstruction^46^ (ASR) based on Riemannian distance calculation for burst removal where the maximum acceptable 0.5-second window standard deviation was set to 100. ASR uses principal components analysis to identify artefact-contaminated, high-amplitude data components, such as those originating from eye blinks or muscle motion, relative to artefact-free reference data. The artefactual signal is rejected and reconstructed in a sliding-window fashion by discarding components with excessive short-window variance (for more details on this procedure, see Mullen et al.^46^). Additionally, problematic data periods were rejected based on the default settings of more than 25% channels showing above-threshold amplitudes (i.e., if ASR did not satisfactorily ‘repair’ data). Data were re-referenced to a common average reference before further data processing.

We performed the extended version of Independent Component Analysis^47^ (ICA) decomposition of input data using the logistic infomax algorithm with 2000 iterations within EEGLab. ICLabel^48^ was used to identify the likely source of each independent component. Components labelled with at least 70% probability of having a non-brain origin (i.e., eye, muscle, heart, line noise, channel noise, other categories) were rejected^49^. For details on the number of rejected components per recording type, see Table S1. This threshold was determined based on a visual inspection of all components derived from the validation dataset, with the intention to discard only components of likely non-brain origin. Finally, a second round of data cleaning was performed using the clean_rawdata plugin with a burst criterion of 20 and all other parameters identical to those in the first round of data cleaning. Pre-processed data were segmented into 2-second epochs. Overall, a similar number of epochs remained following the pre-processing procedure in the pre- and post-task recordings for all datasets (Table S1, Table S2).

### Calculation of periodic and aperiodic EEG metrics

To examine how prominent oscillatory (i.e., iAPF and associated power) and background (i.e., aperiodic exponent) rsEEG metrics associated with age-related changes are affected by cognitive exertion, we first deconstructed the EEG power spectrum into its periodic and aperiodic components. First, we excluded participants from analyses in cases where less than 10 artefact-free epochs remained in the pre- or post-task recordings after pre-processing. Based on this criterion, 24 and 11 participants’s data were excluded from the eyes-open analyses from Session 1 and Session 2, respectively. No participants were excluded in the eyes-closed recordings at either time point. In the validation dataset, 6 participants were excluded from further analyses based on this criterion.

For each channel, time-frequency representations of the epoched data were derived using a multitaper fast Fourier transform with Hanning tapers and a frequency resolution spanning 2 to 40 Hz in steps of 0.2 Hz via the ft_freqanalysis function in FieldTrip. Since narrow-band frequency analyses conflate neural oscillations with broadband background activity, which compromises physiological interpretations^19^, we decomposed the the EEG power spectrum into periodic (alpha peak frequency and related power) and aperiodic components (aperiodic exponent and offset). For each channel, these metrics were iteratively estimated within the frequency range of 2 to 40 Hz^18,22,50^ using the FOOOF algorithm^19^ implemented in FieldTrip. We defined the alpha range from 7 to 14 Hz^22^. All other parameters (including no knee component), were set to the default. In cases where multiple alpha peaks were identified within a given channel, we selected the one with the highest associated power. Channels with a FOOOF model fit below 0.9 were discarded from subsequent analyses^51,52^. For details on the FOOOF model fit related to each recording type, see tables S1 and S2. To prevent confusion stemming from reporting numeric increases or decreases in positive aperiodic exponents versus negative spectral slopes, we adopt the terms ‘steepening’ and ’flattening’ to unambiguously denote directional shifts in the exponent^53^.

Following the above pre-processing procedures, stratification for age and gender (see ‘Participants’ section above) in the DVS dataset was based on data from 595 and 623 participants from the session 1 eyes-open and closed recordings, respectively. Session 2 stratification was employed on data from 317 and 329 participants in the eyes-open and closed conditions, respectively. After pre-processing, the validation dataset included 71 participants (*M* = 69.99 years, *SD* = 3.24 years, 52 females). In further analyses, we utilised spectral metrics derived from a global electrode average. We chose this approach to ensure cross-dataset reproducibility and a topographically unbiased representation of both posterior-dominant alpha and central-parietal aperiodic features.

### Statistical analysis

#### Session 1

We implemented linear mixed-effects modelling within the lme4 package^54^ (version 1.1-37) in R (version 4.5.1; R Core Team, 2025) to evaluate the three EEG metrics of interest (iAPF, power at the iAPF, and the aperiodic exponent) within the Session 1 dataset change as a function of age, sex, task, and eye status. Within the models, we utilised a continuous predictor for age as well as categorical predictors for sex, eye status (i.e., whether data originates from the eyes-open or closed recordings), and task (i.e., pre- or post-task data). We compared 8 different model variants, all of which included an interaction term between eye status and task. Specifically, we compared the following models, defined using standard Wilkinson notation as implemented in the lme4 package^54^;

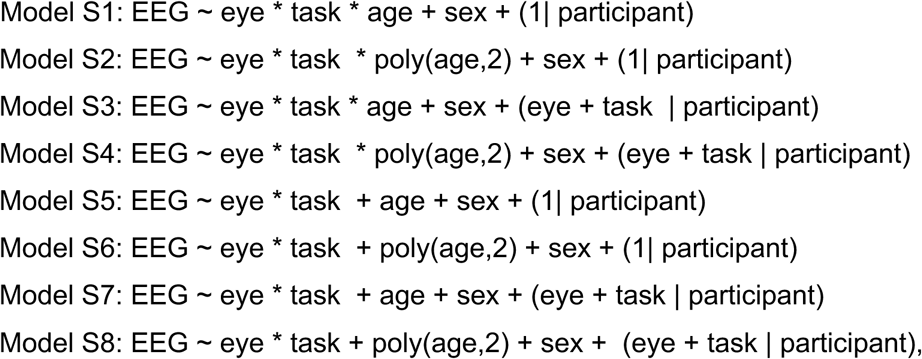

where task refers to data from pre- or post-task recordings and eye indicates data from eyes-open or eye-closed recordings.

To select the best-fitting model for each examined EEG metric, we relied on the Bayesian Information Criterion^55^ (BIC) for model comparison. BICs related to each 8 model variants predicting each EEG metric of interest are shown in Table S3. To correct for multiple comparisons across the three different linear mixed-effects models used to evaluate each of the three EEG measures, we evaluated the significance of the predictors of each winning model against the Bonferroni-corrected *p*-value of 0.0167 (i.e., 0.05/3).

To probe any significant main effects and interactions, we conducted post-hoc analyses using the emmeans package^56^ (version 1.11.2-8) in R (version 4.5.1; R Core Team, 2025). For any significant main effects related to our categorical factors, a post-hoc pairwise comparison was conducted on the estimated marginal means utilising the *‘emtrends*’ function. To characterise the non-linear trajectory of any significant quadratic age effects, a localised post-hoc trend analysis was performed. Because a quadratic model implies that the rate of neural change varies continuously across the lifespan, the instantaneous linear slopes (first derivatives) were estimated at discrete 10-year milestones spanning the middle of age bin (i.e., at the ages of 25, 35, 45, 55, and 65).

To evaluate significant task-age interactions, we employed two different post-hoc testing procedures. First, the full mixed-effects model was sliced at the five discrete 10-year milestones at the middle of each age bin. Within each specific age milestone, the localised conditional marginal means for the pre-task and post-task time points were estimated and contrasted. Next, we calculated and compared the estimated marginal linear slopes of age across the different levels of task to test whether pre- or post-task metrics are more strongly associated with age. For each post-hoc test, we report estimated slopes, its standard errors, *t*-values, Satterthwaite-approximated degrees of freedoms, and *p*-values after multivariate (‘mvt’) adjustment.

As the best-fitting model predicting the aperiodic exponent suggested a stronger effect of age on the post-task compared to pre-task exponents, we utilised Pearson’s correlations to evaluate the strength of these associations. Additionally, we implemented Williams’ *t*-test^57^ via the ‘r_test_paired’ function^58^ in MATLAB (version 2024b, The Mathworks Inc., 2024) to evaluate whether the pre- or post-task EEG exponent is significantly more strongly correlated with age in each eye condition, whilst accounting for the dependence between the two EEG metrics. This approach utilises the correlation matrix among the three variables to derive a *t*-distribution, which tests the null hypothesis that two EEG metrics are equally linearly correlated (two-tailed test) with age when all three sets of observations belong to the same participant (i.e., in a paired observations design).

#### Session 2

To confirm our results in the follow-up Session 2 dataset, we employed the winning model variant for each EEG metrics from Session 1. For iAPF, the model failed to achieve convergence, likely because the reduced sample size was unable to support random slopes alongside a quadratic age predictor. Consequently, we constrained the random effect structure to include random intercepts, but not random slopes, per participant. Model structures for power at iAPF and the aperiodic exponent were identical to those in Session 1. To evaluate significant task-age interactions, the full mixed-effects model was sliced at the five discrete 10-year milestones at the middle of each age bin to derive the localised conditional marginal means for the pre-task and post-task time points. To accommodate the 5-year follow-up interval relative to Session 1, stratum midpoints were centered at ages 30, 40, 50, 60, and 70. Consistent with Session 1, task-dependent differences in age associations were formally evaluated by comparing estimated marginal linear slopes across pre- and post-task recordings. As in Session 1, we employed the same procedure based on Pearson’s correlations and Williams’ *t*-tests to compare the degree of correlation between age and pre-versus post-task exponent.

#### Longitudinal analyses

We employed longitudinal analyses to further confirm our results in the cohort with data from both sessions and evaluate how our EEG metrics change within participants across a 5-year interval. We utilised a similar linear mixed-effects model procedure as above, which also accounted for session type. For each EEG metrics of interest, we utilised the BIC to select the best-fitting model from following model variants;

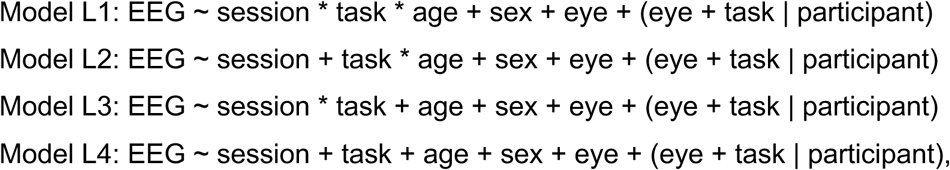

where session is a categorical predictor referring to Session 1 or 2. To prevent model over-parameterisation and avoid drastically expanding the model matrix over a 4-way interaction (i.e., eye x session x task x age), candidate models were constrained to a maximum of a three-way interaction term (i.e., session x task x age), with eye condition modelled as a fixed effect. As model selection in Session 1 established optimal fits of models including random slopes and intercepts, we maintained this random-effects specification across longitudinal candidate models. BICs related to each 4 model variants predicting each EEG metric are shown in Table S5. Significant effects within each winning model were evaluated using the same post-hoc testing procedures as in the Session 1 analyses.

#### Validation dataset

We employed and compared the following two linear mixed-effect model variants;

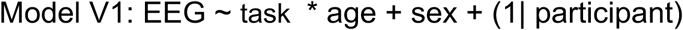

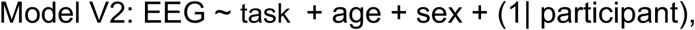

where task refers to whether data originates from the pre- or post-task recording. Because models including random slopes failed to converge on this dataset, random effects were constrained to random intercepts per participant. BICs related to the 2 model variants predicting each EEG are shown in Table S6.

#### Model fit

To evaluate the fit of each winning model, we calculated marginal (*R^2^*m) and conditional (*R^2^c*) coefficients of determination (*R^2^*). The former represents the proportion of total variance explained by fixed effects alone, whereas the latter reflects the proportion of variance explained by the combination of fixed and random effects. Computations were implemented in the performance package^59^ (version 0.17.1) in R (version 4.5.1; R Core Team, 2025).

## Results

### Age- and Task-Related Changes in EEG frequency metrics (Session 1)

To investigate how prominent age-related rsEEG markers (iAPF, its associated power, and the aperiodic exponent) respond to acute cognitive exertion across the adult lifespan, we evaluated 8 candidate models for each rsEEG metric (Table S2). Table 1 displays the means and standard deviations related to each spectral metric separately for each recording type. Grand-average power spectra and scalp topographies for each rsEEG metric, shown separately for pre- and post-task recordings, are illustrated in Fig. 2. For an overview of parameter estimates and associated *p*-values from the winning model predicting each rsEEG metric, see Table 2.

**Figure 2.**
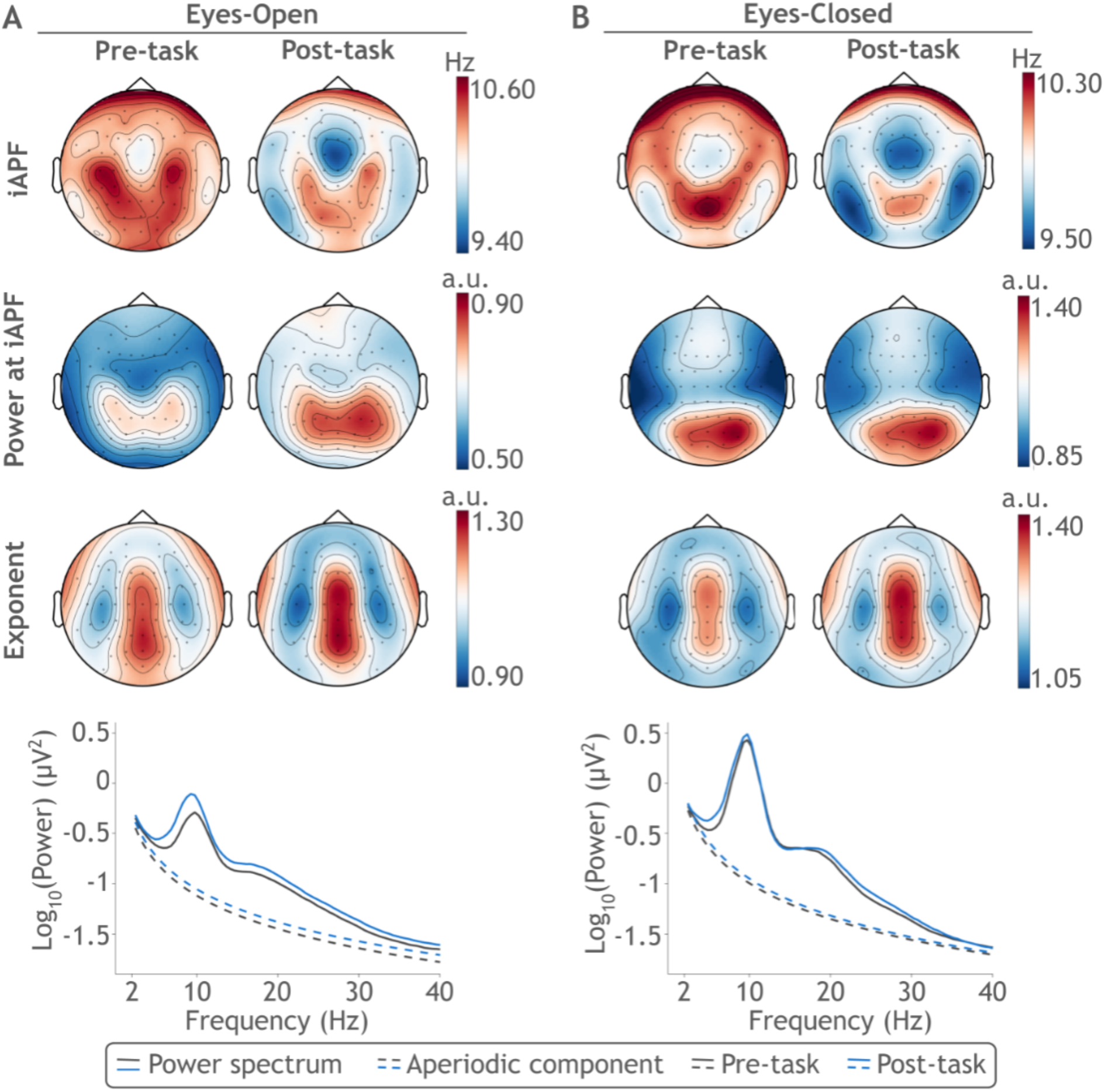
Scalp topographies and power spectra before and after the cognitive task battery in Session 1 of the Dortmund Vital Study, averaged for the eyes open (A) and closed (B) conditions. In each panel, the left and right side shows pre- and post-task topographies, respectively, averaged across participants. Horizontally, the topographies depict individual alpha peak frequency (iAPF; top), its associated power (middle), and the aperiodic exponent (bottom). The bottom plot of each panel shows the mean raw power spectra (continuous lines) and aperiodic components (dashed lines) within the respective eye conditions, compared between pre-task (grey) and post-task (blue) recordings, and averaged across all channels and participants.

**Table 1.** Descriptive statistics for spectral metrics in Session 1 (N = 390). Within each cell, the mean (standard deviation) is shown for individual alpha peak frequency (iAPF), power at iAPF, and the aperiodic exponent, separately for eye condition (eyes-open vs. eyes closed) and task time point (pre-task vs. post-task).

|  |  | iAPF<br>(Hz) | Power<br>at iAPF (a.u.) | Aperiodic<br>Exponent<br>(a.u.) |
| --- | --- | --- | --- | --- |
| Eyes- Open | Pre-task | 10.32<br>(1.01) | 0.60<br>(0.31) | 1.10<br>(0.20) |
|  | Post-task | 10.04<br>(1.00) | 0.69<br>(0.32) | 1.09<br>(0.23) |
| Eyes- Closed | Pre-task | 10.05<br>(0.93) | 1.06<br>(0.43) | 1.18<br>(0.22) |
|  | Post-task | 9.82<br>(0.85) | 1.07<br>(0.42) | 1.23<br>(0.25) |

**Table 2.** Parameter estimates and associated *p*-values for the winning linear mixed-effects models predicting resting-state EEG metrics in Session 1. Coefficients (*b*) and *p*-values (*p*) are presented for individual alpha peak frequency (iAPF), power at iAPF, and the aperiodic exponent. Model variants were evaluated using the Bayesian Information Criterion (BIC), with full model comparison details provided in Table S3. Reference levels for categorical predictors are pre-task for task (pre-task vs. post-task), closed for eye condition (closed vs. open), and fe*male* for sex (female vs. male). Age was a continuous predictor between 20 and 70 years. Predictors were evaluated against a Bonferroni-corrected significance threshold of *p* < 0.0167.

|  | iAPF |  | Power at iAPF |  | Aperiodic Exponent |  |
| --- | --- | --- | --- | --- | --- | --- |
| | $b$ | $p$ | $b$ | $p$ | $b$ | $p$ |
| Intercept | 10.47 | <0.001 | 0.87 | <0.001 | 1.31 | <0.001 |
| Eye | -0.28 | <0.001 | 0.46 | <0.001 | 0.12 | <0.001 |
| Task | -0.29 | <0.001 | 0.09 | <0.001 | 0.13 | <0.001 |
| Age (1st Order) | -4.57 | 0.004 | -0.01 | <0.001 | -0.01 | <0.001 |
| Age (2nd Order) | -5.51 | <0.001 |  |  |  |  |
| Sex | -0.30 | <0.001 | -0.01 | 0.790 | 0.03 | 0.049 |
| Eye x Task | 0.07 | 0.109 | -0.08 | <0.001 | 0.02 | 0.350 |
| Eye x Age |  |  |  |  | -0.0007 | 0.160 |
| Task x Age |  |  |  |  | -0.003 | <0.001 |
| Eye x Task x Age |  |  |  |  | 0.0005 | 0.305 |

#### iAPF

For iAPF, Model S8 (*R^2^m* = 0.09, *R^2^c* = 0.83) was selected as the optimal model (Table S3) as it yielded the lowest BIC value, indicating the best balance between model fit and complexity^55^. This model revealed that iAPF was significantly higher pre-compared to the post-task (*b* = -0.29, *SE* = 0.03., *t*(762) = -9.21, *p* < 0.001), in the eyes-open compared to eyes-closed recordings (*b* = -0.28, *SE* = 0.04, *t*(633) = -6.98, *p* < 0.001), and in females compared to males (*b* = -0.30, *SE* = 0.08, *t*(390) = -3.77, *p* < 0.001). There was no significant interaction between eye status and task (*b* = 0.07, *SE* = 0.04, *t*(390) = 1.60, *p* = 0.11).

There was a significant negative linear trend for age (*b* = -4.57, *SE* = 1.57, *t*(390) = -2.92, *p* = 0.004), which was qualified by a significant negative quadratic age trend (*b* = -5.51, *SE* = 1.57, *t*(390) = -3.52, *p* < 0.001). Together, this implies that the trajectory of iAPF followed a concave path where the rate of decline accelerated significantly in older cohorts (Fig. 3). Post-hoc analysis revealed that the instantaneous linear slope associated with age was significantly positive at age 25 (slope = 0.02, *SE* = 0.009, *t(*394) = 2.46, *p* = 0.04), but not at age 35 (slope = 0.07, *SE* = 0.005, *t(*394) = 1.40, *p* = 0.35). Then this slope became significantly negative and progressively steeper; accelerating from age 45 (slope = -0.009, *SE* = 0.003, *t(*394) = -3.04, *p* = 0.009), through age 55 (slope = -0.02, *SE* = 0.005, *t*(394) = -4.47, *p* < 0.001) and 65 (slope = -0.04, *SE* = 0.01, *t*(394) = -4.17, *p* < 0.001).

**Figure 3.**
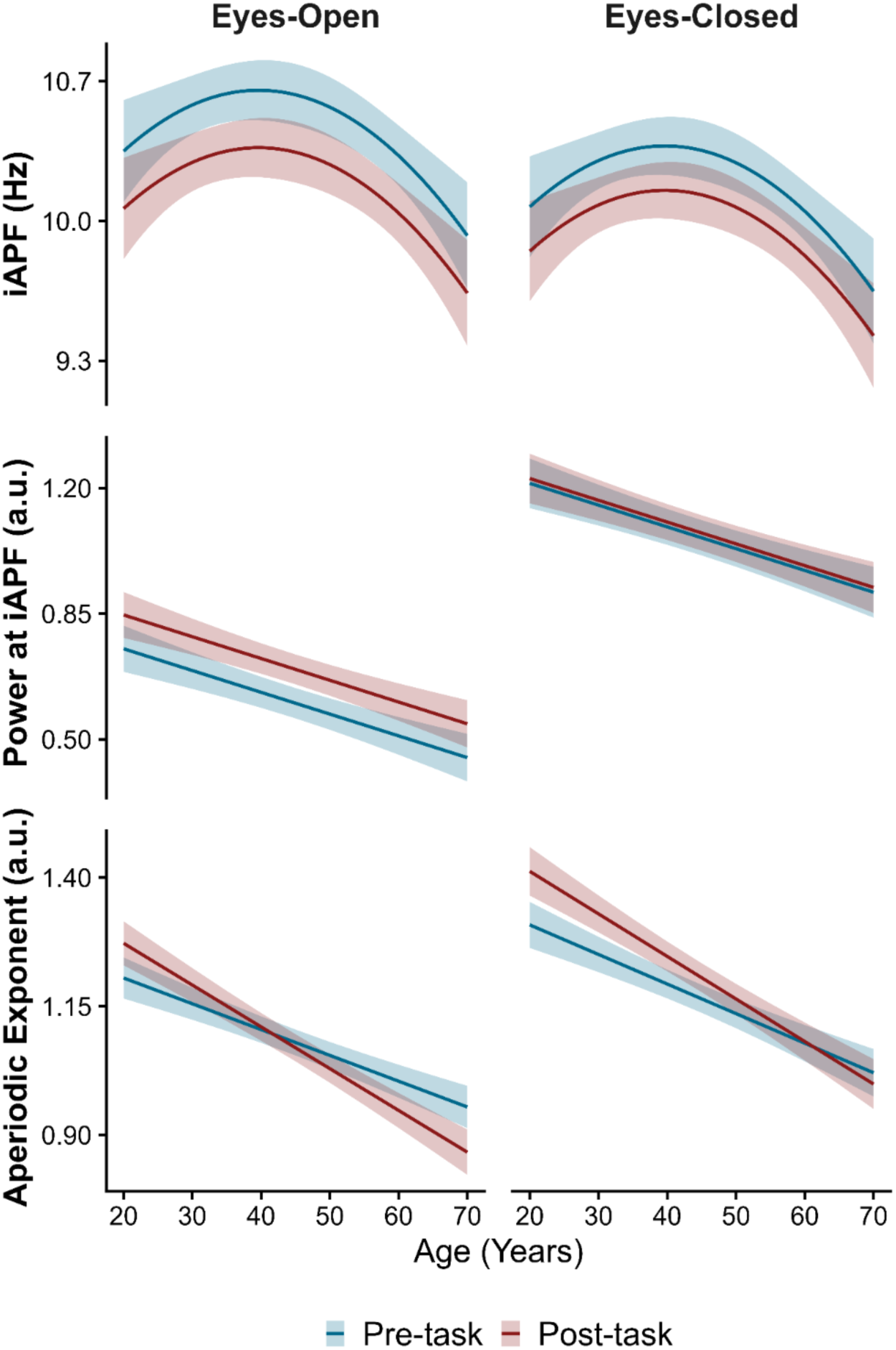
Predicted lifespan trajectories of periodic and aperiodic rsEEG markers in Session 1. Trajectories are based on the winning linear mixed-effects model for each EEG metric (Table S2 Table 1), representing the predicted marginal effects of chronological age (x-axis) on the electrophysiological markers (y-axis). The continuous lines and color-matched shaded ribbons depict these predictions and their 95% confidence intervals, comparing pre-task (blue) and post-task (red) recordings. The left and right columns represent predictions for eyes-open and eyes-closed recordings, respectively. Vertically, the panels depict trajectories for individual alpha peak frequency (iAPF; top), its associated power (middle), and the aperiodic exponent (bottom).

#### Power at the iAPF

Power at the iAPF was best explained by Model S7 (*R^2^m* = 0.29, *R^2^c* = 0.89; Table S3). We found a significant eye-by-task interaction (*b* = -0.08, *SE* = 0.01, *t*(390) = -5.55, *p* < 0.001; Fig. 3), which superseded the significant main effect of eye status (*b* = 0.46, *SE* = 0.02, *t*(567) = 27.85, *p* < 0.001) as well as the non-significant main effect of task (*b* = 0.09, *SE* = 0.01, *t*(773) = 8.70, *p* = 0.22). We utilised pairwise post-hoc contrasts to evaluate the impact of task within each eye state as well as eye state pre- and post-task. The eyes-open condition was associated with a significant increase in power from pre- to post-task (pre-post contrast estimate = -0.09, SE = .01, *t(*775) = -8.69, *p* < 0.001). For the eyes-closed condition, power remained stable between pre- and post-task (contrast estimate = -0.01, SE = 0.01, *t*(775) = -1.24, *p* = 0.22). Second, power was significantly higher in the eyes-closed condition relative to the eyes-open condition both pre- (open-closed contrast estimate = -0.46, SE = 0.02, *t*(568) = -27.81, *p* <0 .001) and post-task (open-closed contrast estimate = -0.38, SE = 0.02, *t(*568) = -22.93, *p* < 0.001). Together, these contrasts confirm that the eye-by-time interaction in our model was driven by a task effect that is restricted to the open-eye state, which also narrowed the discrepancy between the two eye conditions post-task. Regarding demographic variables, there was a significant negative linear main effect of age (*b* = -0.006, *SE* = 0.001, *t(*390) = -5.96, *p* < 0.001) but the effect of sex was non-significant (*b* = -0.008, *SE* = 0.03, *t*(390) = -0.27, *p* = 0.79).

#### Aperiodic Exponent

The aperiodic exponent was best explained by Model S3 (*R^2^m* = 0.24, *R^2^c* = 0.88; Table S3). Crucially, a significant two-way interaction emerged between task and age (*b* = - 0.003, *SE* = 0.0005, *t*(703) = -6.30, *p* < 0.001; Fig. 3), which superseded the standalone main effects of age (*b* = -0.005, *SE* = .0006, *t*(430) = -7.74, *p* < 0.001) and task (*b* = 0.13, *SE* = .02, *t*(703) = 5.61, *p* < 0.001). The three-way interaction between eye status, time, and age was not statistically significant *(b* = 0.0006, *SE* = 0.0006, *t*(390) = 1.03, *p* = 0.31). Similarly, the two-way interactions between eye status and task (*b* = 0.02, *SE* = 0.03, *t*(390) = 0.94, *p* = 0.35) and eye status and age (*b* = -0.0007, *SE* = 0.0005, *t*(672) = -1.41, *p* = 0.16) were not significant. We found a significant main effect of eye status (*b* = 0.12, *SE* = 0.02, *t*(672) = 4.83, *p* < 0.001), indicating overall flatter exponents in the eyes-open recordings. The main effect of sex (*b* = 0.03, *SE* = 0.02, *t*(390) = 1.97, *p* = 0.05) was not significant when evaluated against the Bonferroni-corrected *p*-value of 0.0167.

To unpack the interaction between task and age, we first evaluated the simple main effects of the task condition at 10-year milestones ranging from age 25 to 65. This analysis revealed a developmental crossover effect between task time points. For younger cohorts, the task elicited a significant increase in the aperiodic exponent, reaching its maximum shift at age 25 (pre-post contrast estimate = -0.07, *SE* = 0.01, *t*(392) = -7.33, *p* < 0.001), and remaining significant through ages 35 (pre-post contrast estimate = -0.04, *SE* = 0.007, *t*(392) = -6.27, *p* < 0.001) and 45 (pre-post contrast estimate = -0.02, *SE* = 0.006, *t*(392) = -2.67, *p* = 0.008). At 55 years, there was no task-induced change between pre- and post-task recordings (pre-post contrast estimate = 0.01, *SE* = 0.007, *t*(392) = 1.78, *p* = 0.08). Crucially, in older cohorts, this pattern significantly reversed. We observed a significant flattening of the exponent from pre- to post-task at age 65 (pre-post contrast estimate = 0.04, *SE* = 0.01, *t*(392) = 4.06, *p* < 0.001).

To test whether the effect of age is stronger in pre- or post-task exponents, we compared the continuous linear slopes of age pre- and post-task. The exponent was significantly and negatively associated with age both before (*b* = -0.005, *SE* = 0.0006, t(393) = -8.30, *p* < 0.001) and after the task (*b* = -0.008, *SE* = 0.0007, *t*(393) = -11.66, *p* < .001). Furthermore, the estimated marginal slope of age was significantly more negative in post-task recordings (pre-post slope difference = 0.003, SE = 0.0004, t(392) = 6.97, p < 0.001), suggesting a stronger age effect post-task. In alignment with the modelling results, Pearson’s correlations indicated a significant correlation between age and both the pre-task (eyes-open: *r* = -0.36, *p* < 0.001, eyes-closed: *r* = -0.37, *p* < 0.001) and post-task (eyes-open: *r* = -0.50, *p* < 0.001, eyes-closed: *r* = -0.47, *p* < 0.001) exponents. Crucially, Williams’ *t*-tests^57^ suggested that compared to pre-task, post-task exponents were significantly more strongly associated with age in both eyes-open (*p* < 0.001, *t*(387) = 4.52) and the eyes-closed (*p* < 0.001, *t*(387) = 3.77) recordings.

### Validation within the Session 2 dataset

To assess the reliability of the Session 2 data, we computed intra-class correlation coefficients measuring absolute agreement^60^ (ICC) between Session 1 and 2 spectral metrics for each eye condition and time point (pre-task, post-task). Similarly to previous research utilising the longitudinal sub-sample of the DVS^7^, all spectral measures were characterised by at least moderate reliability (Table S4). To validate the age and task-related effects observed in the Session 1 data, we applied the best-fitting models for each EEG metric from Session 1 to the Session 2 dataset. Pre- and post-task grand-average power spectra and scalp topographies for each rsEEG metric are shown in Fig. 4. Table 3 provides an overview of parameter estimates and associated *p*-values for the linear-mixed effect model predicting each of the spectral measures. Table 4 displays the means and standard deviations related to each spectral metric separately for each recording type.

**Figure 4.**
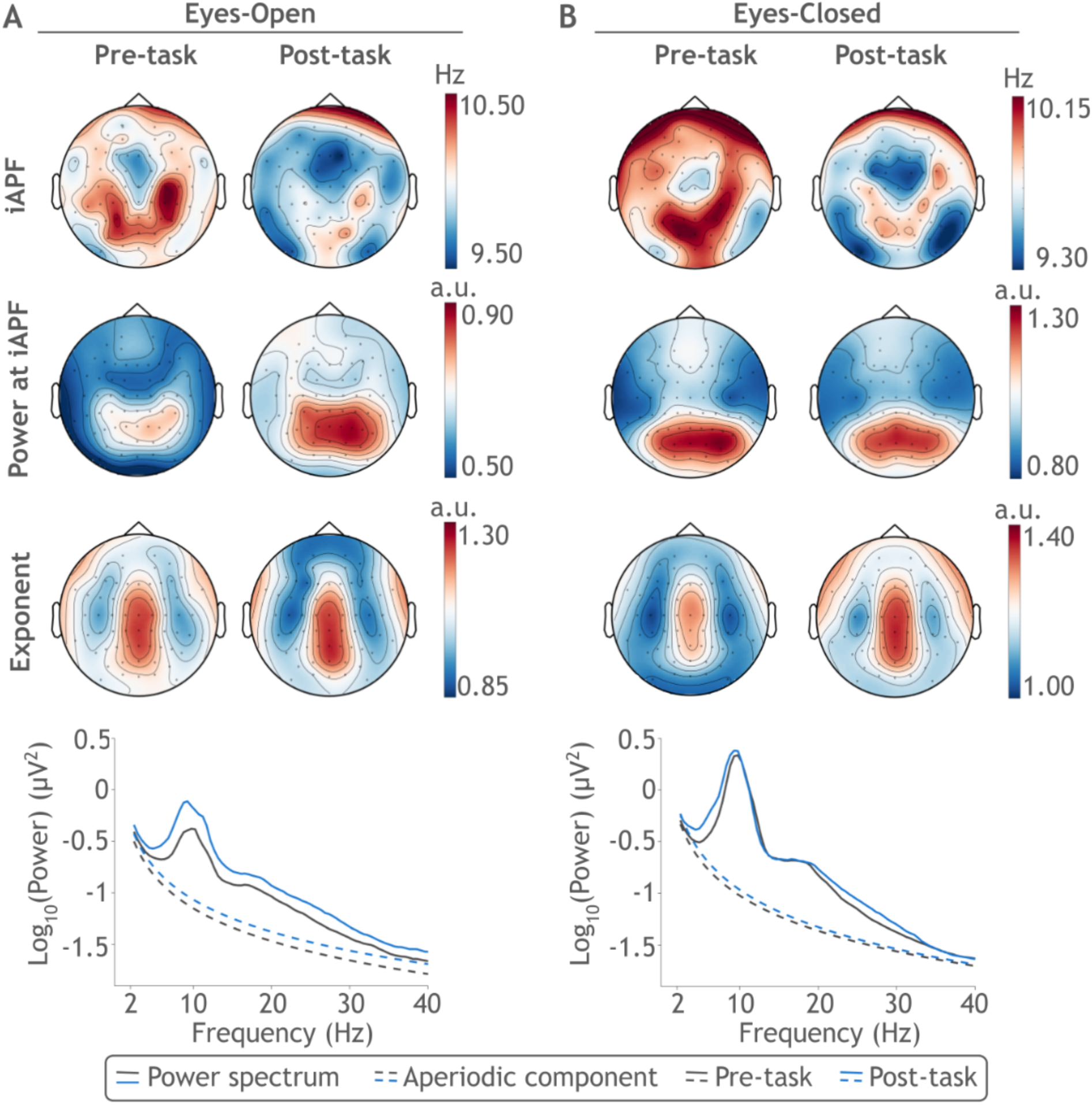
Scalp topographies and power spectra before and after the cognitive task battery in Session 2 of the Dortmund Vital Study, averaged separately for the eyes open (A) and closed (B) conditions. In each panel, the left and right side shows pre- and post-task topographies, respectively, averaged across participants. Horizontally, the topographies depict individual alpha peak frequency (iAPF; top), its associated power (middle), and the aperiodic exponent (bottom). The bottom plot of each panel shows the mean raw power spectra (continuous lines) and aperiodic components (dashed lines) within the respective eye conditions, compared between pre-task (grey) and post-task (blue) recordings, and averaged across all channels and participants.

**Table 3.**
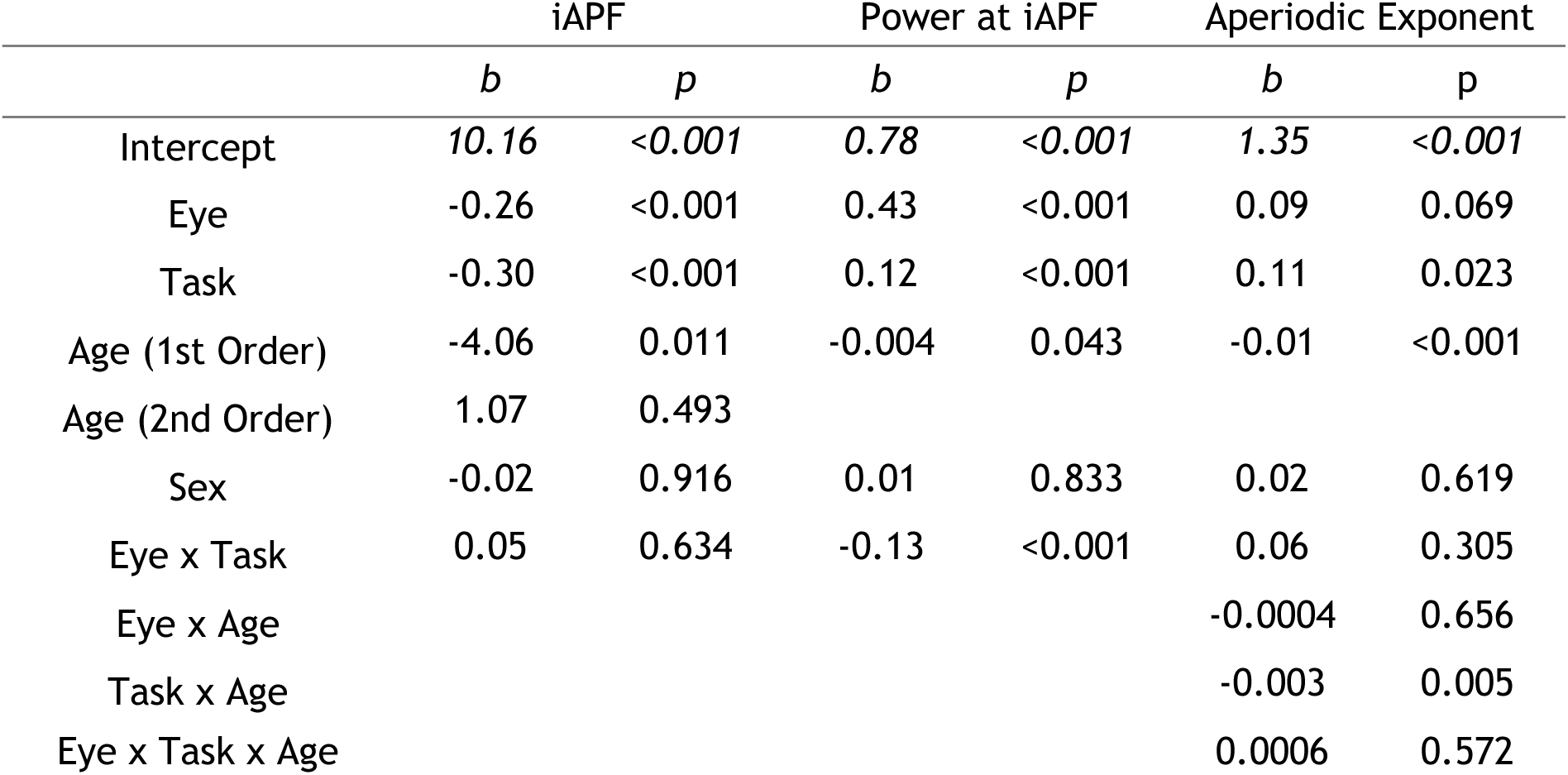
Parameter estimates and associated *p*-values for the linear mixed-effects models predicting resting-state EEG metrics in Session 2. Coefficients (*b*) and *p*-values (*p*) are presented for individual alpha peak frequency (iAPF), power at iAPF, and the aperiodic exponent. Models were fitted using the optimal winning model established in Session 1. As this winning model failed to converge for iAPF, results are reported from Model S6, which no longer contains participant-wise random slopes for eye and task condition. Reference levels for categorical predictors are pre-task for task (pre-task vs. post-task), closed for eye condition (closed vs. open), and female for sex (female vs. male). Age was a continuous predictor between 25 and 75 years. Statistically significant predictors (*p* < 0.05) highlight the replication of key Session 1 effects, notably the task-by-age interaction for the aperiodic exponent.

|  | iAPF |  | Power at iAPF |  | Aperiodic Exponent |  |
| --- | --- | --- | --- | --- | --- | --- |
| | $b$ | $p$ | $b$ | $p$ | $b$ | $p$ |
| Intercept | 10.16 | <0.001 | 0.78 | <0.001 | 1.35 | <0.001 |
| Eye | -0.26 | <0.001 | 0.43 | <0.001 | 0.09 | 0.069 |
| Task | -0.30 | <0.001 | 0.12 | <0.001 | 0.11 | 0.023 |
| Age (1st Order) | -4.06 | 0.011 | -0.004 | 0.043 | -0.01 | <0.001 |
| Age (2nd Order) | 1.07 | 0.493 |  |  |  |  |
| Sex | -0.02 | 0.916 | 0.01 | 0.833 | 0.02 | 0.619 |
| Eye x Task | 0.05 | 0.634 | -0.13 | <0.001 | 0.06 | 0.305 |
| Eye x Age |  |  |  |  | -0.0004 | 0.656 |
| Task x Age |  |  |  |  | -0.003 | 0.005 |
| Eye x Task x Age |  |  |  |  | 0.0006 | 0.572 |

**Table 4.**
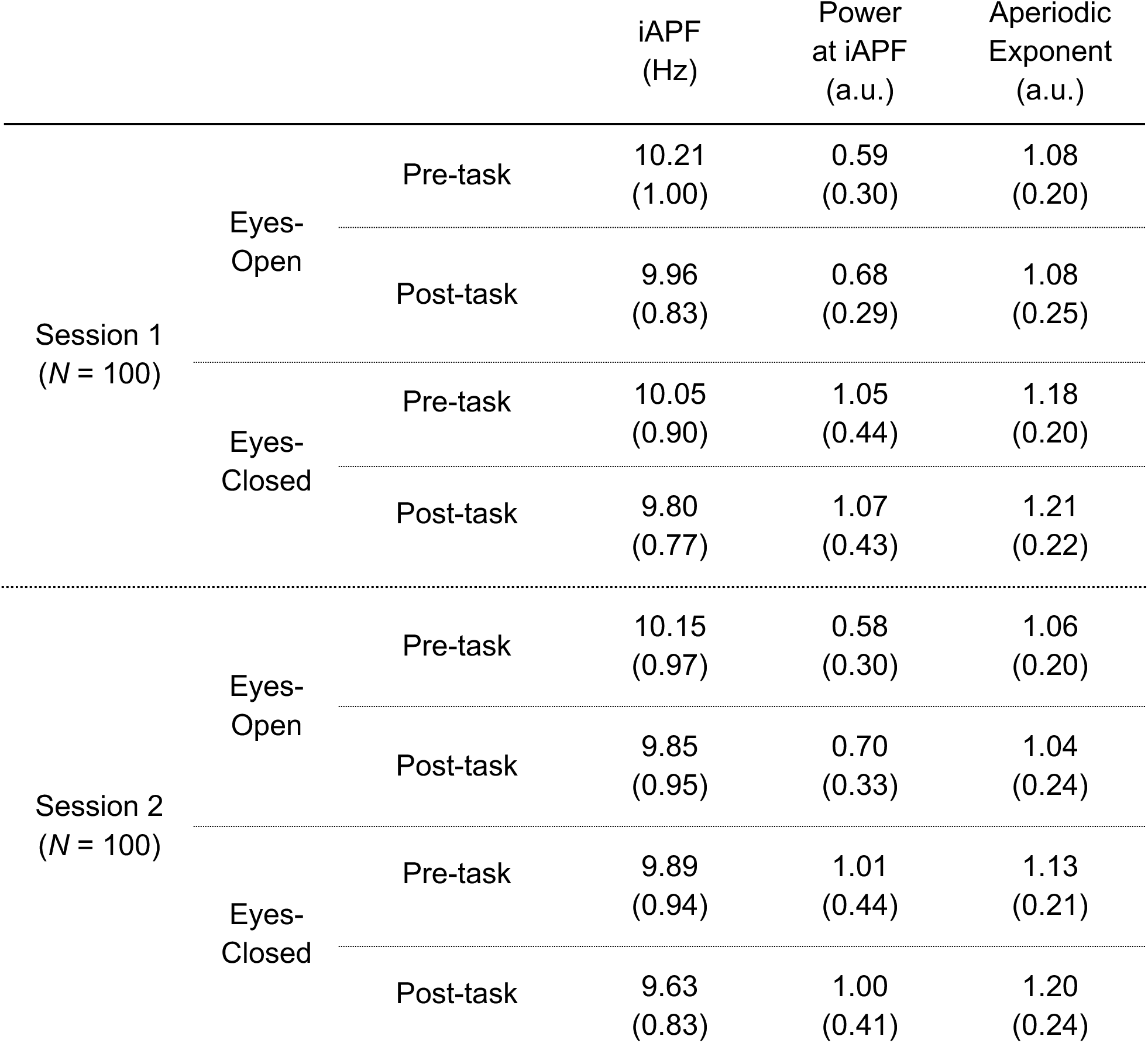
Descriptive statistics for spectral metrics for the longitudinal sub-sample (*N* = 100). Within each cell, the mean (standard deviation) is shown for individual alpha peak frequency (iAPF), power at iAPF, and the aperiodic exponent, separately for Session (Session 1 vs Session2), eye condition (eyes-open vs. eyes closed), and task time point (pre-task vs. post-task).

#### iAPF

As the best-fitting Session 1 model for iAPF (Model S8) failed to converge when fitted to Session 2 data, we simplified the random-effects structure to include random intercepts, but not random slopes, per participant (i.e., Model S6; *R^2^m* = 0.08, *R^2^c* = 0.66). Similarly to the Session 1 results, this model revealed significantly lower iAPF in the post-task compared to pre-task recordings (*b* = -0.30, SE = 0.07, *t*(300) = 4.19, *p <* 0.001; Fig. 5) as well as in the eyes-closed compared to the eye-open condition (*b* = -0.26, SE = .07, *t*(300) = -3.69, *p <* 0.001), without a significant interaction between eye status and task (*b* = 0.05, SE = 0.10, *t*(300) = 0.48, *p* = 0.63). Whilst the negative linear relationship between age and iAPF (*b* = -4.06, SE = 1.56, *t*(100) = -2.60, *p* = 0.01) was replicated, this was not qualified by a significant quadratic age trend (*b* = 1.07, SE = 1.56, *t*(100) = -0.69, *p* = 0.49). Further in contrast with the Session 1 results, the effect of sex was not significant (*b* = -0.02, SE = 0.16, *t*(100) = -0.11, *p* = 0.92).

**Figure 5.**
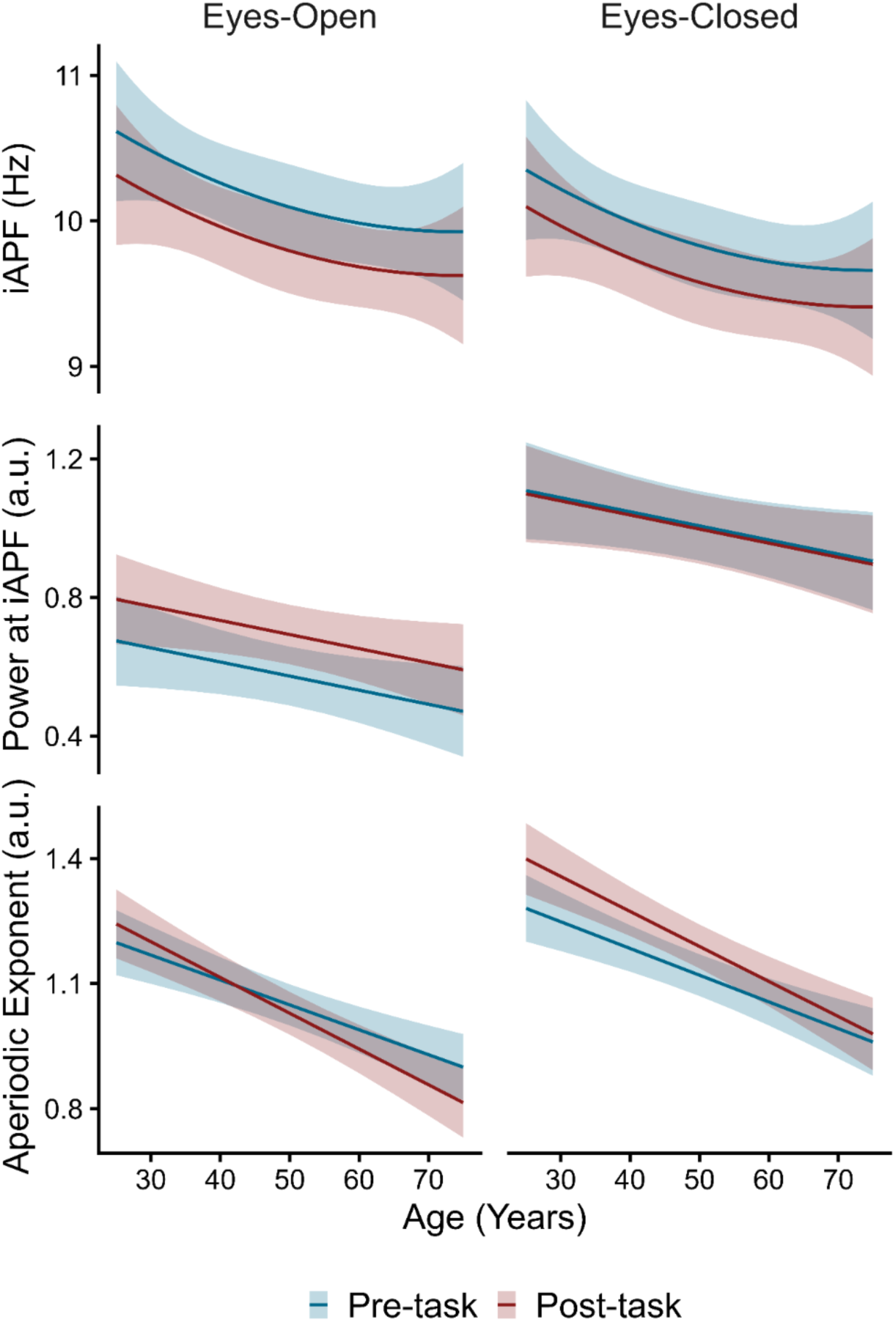
Predicted lifespan trajectories of periodic and aperiodic rsEEG markers in Session 2. Trajectories are based on the winning linear mixed-effects model structures established in Session 1, fitted to the Session 2 data to represent the predicted marginal effects of chronological age (x-axis) on the electrophysiological markers (y-axis). For iAPF, the model was simplified to exclude random slopes. The continuous lines and color-matched shaded ribbons depict these predictions and their 95% confidence intervals, comparing pre-task (blue) and post-task (red) recordings. The left and right columns represent predictions for eyes-open and eyes-closed recordings, respectively. Vertically, the panels depict trajectories for individual alpha peak frequency (iAPF; top), its associated power (middle), and the aperiodic exponent (bottom).

To determine whether the discrepancies between the Session 1 and 2 results stem from reduced statistical power (*N* = 390 in Session 1, *N* = 100 in Session 2) or longitudinal shifts over the 5-year interval, we fitted Model 6 to the Session 1 data restricted to the sub-set of 100 participants included in Session 2. Within this Session 1 sub-sample, the pattern of results mirrored those from Session 2. Whilst the negative linear age effect remained statistically significant (*b* = -3.92, SE = 1.41, *t*(100) = 2.78, *p =* 0.006), the quadratic age component was not (*b* = -1.26, SE = 1.41, *t*(100) = -0.89, *p =* 0.37). The main effect of sex (*b* = 0.04, SE = 0.14, *t*(100) = 0.05, *p =* 0.96) and eye status were non-significant (*b* = -0.15, SE = 0.08, *t*(300) = -1.96, *p = 0.051*), although the latter approached statistical significance. Task effects remained robust (Fig. S1) as iAPF was significantly lower post-task compared to pre-task (*b* = -0.25, SE = 0.08, *t*(300) = -3.23, *p = 0.001*), with no significant eye-by-task interaction (*b* = -0.003, SE = .11, *t*(300) = -0.03, *p = 0.98*). These results indicate that the absence of a significant quadratic age trend and sex effect in Session 2 is attributable to lower statistical power in the sub-sample rather than longitudinal changes over time.

#### Power at the iAPF

Model S7 (*R^2^m* = 0.24, *R^2^c* = 0.85) revealed identical effects related to power at the iAPF as in Session 1. Specifically, we found a significant negative main effect of age (*b* = - 0.004, *SE* = 0.002, *t*(100) = -2.03, *p* = 0.045), but not of sex (*b* = 0.01, *SE* = 0.06, *t*(100) = 0.21, *p* = 0.83). Further in accordance with the Session 1 results, there was a significant interaction between eye status and task (*b* = -0.13, *SE* = 0.03, *t*(100) = -4.48, *p* < 0.001; Fig. 5), which superseded the significant main effects of eye (*b* = 0.43, *SE* = 0.03, *t*(154) = 14.33, *p* < 0.001) and task (*b* = 0.12, *SE* = 0.02, *t*(197) = 5.48, *p* < 0.001).

Pairwise post-hoc tests revealed the same effects as observed in Session 1. Firstly, power at the iAPF was significantly higher in the eyes-closed compared to the eyes-open recordings both pre- (open-closed contrast estimate = -0.43, *SE* = 0.03, t(156) = -14.25, *p* < 0.001) and post-task (open-closed contrast estimate = -0.31, *SE* = 0.03, t(156) = 10.01, *p* < 0.001). Secondly, power at the iAPF significantly increased from pre- to post-task in the eyes-open (pre-post contrast estimate = -0.12, *SE* = 0.02, *t(*199) = -5.45, *p* < 0.001), but not in the eyes-closed recordings (pre-post contrast estimate = 0.01, *SE* = 0.02, *t(*199) = 0.42, *p* = 0.67).

#### Aperiodic Exponent

Model S3 (*R^2^m* = 0.31, *R^2^c* = 0.81) revealed a very similar pattern of results on the aperiodic exponent in Sessions 1 and 2. Firstly, the three-way interaction between time, eye conditions, and age (*b* = 0.0006, *SE* = 0.001, *t*(100) = 0.57, *p* = 0.57), the two-way interactions between eye status and task (*b* = 0.06, *SE* = 0.06, *t*(100) = 1.03, *p* = 0.31), and the two-way interaction between eye status and age (*b* = -0.0004, *SE* = 0.001, *t*(174) = -0.45, p = 0.66) were not statistically significant. The effect of eye condition (*b* = 0.09, *SE* = 0.05, *t*(174) = 1.82, *p* = 0.07) and sex was not significant (*b* = 0.02, *SE* = 0.04, *t*(100) = 0.50, *p* = 0.62). Crucially, the significant two-way interaction between task and age (*b* = -0.002, *SE* = 0.001, *t*(182) = -2.80, *p* = 0.006; Fig. 5) was replicated, which superseded the standalone main effects of age (*b* = -0.006, *SE* = 0.001, *t*(109) = -4.79, *p* < 0.001) and task (*b* = 0.11, *SE* = 0.05, *t*(182) = 2.28, *p* = 0.02).

Next, we evaluated the simple main effects of the recording time at 10-year lifespan milestones at the middle of each age stratum (i.e., at ages 30, 40, 50, 60, 70), which were shifted by 5 five years compared to Session 1. As in Session 1, the task elicited a significant increase in the aperiodic exponent for younger cohorts at the ages of 30 (pre-post contrast estimate = -0.07, *SE* = 0.02, *t*(102) = -3.73, *p* < 0.001), 40 (pre-post contrast estimate = -0.05, *SE* = 0.01, *t*(102) = -3.50, *p* < 0.001), and 50 (pre-post contrast estimate = -0.01, *SE* = 0.01, *t*(102) = -2.16, *p* = 0.03) but not at the ages of 60 (pre-post contrast estimate = -0.001, *SE* = 0.01, *t*(102) = -0.11, *p* = 0.91) and 70 (pre-post contrast estimate = 0.02, *SE* = 0.02, *t*(102) = 1.13, *p* = 0.26). As in Session 1, post-hoc tests revealed that whilst both the pre- (*b* = -0.006, *SE* = 0.001, *t*(103) = -5.24, *p* < 0.001) and post-task (*b* = -0.008, *SE* = 0.001, *t*(103) = -6.56, *p* < 0.001) exponents were negatively related to age, there was a stronger effect of age on the post-task compared to the pre-task exponent (slope difference = -0.002, *SE* = 0.0008, *t*(102) = -3.01, *p* = 0.003).

Similarly to Session 1, Pearson’s correlations indicated a significant correlation between age and both the pre-task (eyes-open: *r* = -0.45, *p* < 0.001, eyes-closed: *r* = -0.45, *p* < 0.001) and post-task (eyes-open: *r* = -0.52, *p* < 0.001, eyes-closed: *r* = -0.49, *p* < 0.001) exponents. However, Williams’ *t*-tests revealed no significant difference in the strength of the pre-versus post-task associations in either the eyes-open (*p* = 0.17, *t*(97) = 1.39) or eyes-closed (*p* = 0.17, *t*(97) = 1.38) conditions. Despite similar descriptive trends toward stronger age correlations post-task, the reduced sample size in Session 2 likely limited the statistical power to detect significant differences.

### Longitudinal analysis

We leveraged our longitudinal sub-sample of participants with data from both sessions to test whether the EEG metrics exhibit true within-person longitudinal change over the 5-year period and whether task-related dynamics observed cross-sectionally are reliably preserved within the same individuals across the two sessions. Table 4 displays the means and standard deviations related to each spectral metric separately for each recording type. For each EEG metrics of interest, we tested four linear mixed-effects model variants and evaluated the winning model selected based on the lowest BIC (Table S5). Table 5 contains an overview of parameter estimates and associated *p*-values for each winning model.

**Table 5.** Parameter estimates and associated *p*-values for the winning linear mixed-effects models predicting longitudinal resting-state EEG metrics across Sessions 1 and 2. Coefficients (*b*) and *p*-values (*p*) are presented for individual alpha peak frequency (iAPF), power at iAPF, and the aperiodic exponent. Optimal model structures were selected based on the Bayesian Information Criterion (BIC), with full model comparison details provided in Table S5. Reference levels for categorical predictors are Session 1 for Session (Session 1 vs. Session 2), pre-task for task (post-task vs. pre-task), closed for eye condition (closed vs. open), and female for sex (female vs. male). Continuous age was measured across the lifespan between the ages of 20 and 70. We observed significant within-person longitudinal declines over the 5-year follow-up (Session) in the iAPF and the aperiodic exponent as well as a preserved task-by-age interaction for the aperiodic exponent.

|  | iAPF |  | Power at iAPF |  | Aperiodic Exponent |  |
| --- | --- | --- | --- | --- | --- | --- |
| | $b$ | $p$ | $b$ | $p$ | $b$ | $p$ |
| Intercept | 10.81 | <0.001 | 0.80 | <0.001 | 1.35 | <0.001 |
| Session | -0.12 | 0.004 | -0.02 | 0.120 | -0.03 | 0.005 |
| Task | -0.26 | <0.001 | 0.06 | <0.001 | 0.13 | <0.001 |
| Age | -0.01 | 0.006 | -0.003 | 0.062 | -0.01 | <0.001 |
| Sex | 0.003 | 0.985 | -0.05 | 0.406 | 0.01 | 0.848 |
| Eye | -0.20 | <0.001 | 0.40 | <0.001 | 0.11 | <0.001 |
| Task x Age |  |  |  |  | -0.002 | <0.001 |

#### iAPF

For iAPF, the best-fitting model (Model L4; *R^2^m* = 0.08, *R^2^c* = 0.79; Table S5) included main effects for session, task, eye condition, age, and sex, without any interaction between predictors. In addition to the significant cross-sectional decrease in iAPF as a function of age (*b* = -0.01, *SE* = 0.005, *t*(100) = -2.78, *p* = 0.006), we found that the iAPF also significantly decreased within-participants over the 5-year period (main effect of session: *b* = -0.12, *SE* = 0.04, *t*(100) = -2.93, *p* = 0.004). Similarly to the Session 1 and 2 results, we found that iAPF was significantly higher pre-compared to post-task (*b* = -0.26, *SE* = 0.04, *t*(100) = -6.46, *p* < 0.001; Fig. 6) as well as in the eyes-open compared to the eyes-closed condition (*b* = -0.20, *SE* = 0.06, *t*(100) = -3.51 *p* < 0.001). There was no significant effect of sex (*b* = 0.003, *SE* = 0.06, *t*(100) = 0.02, *p* = 0.99).

**Figure 6.**
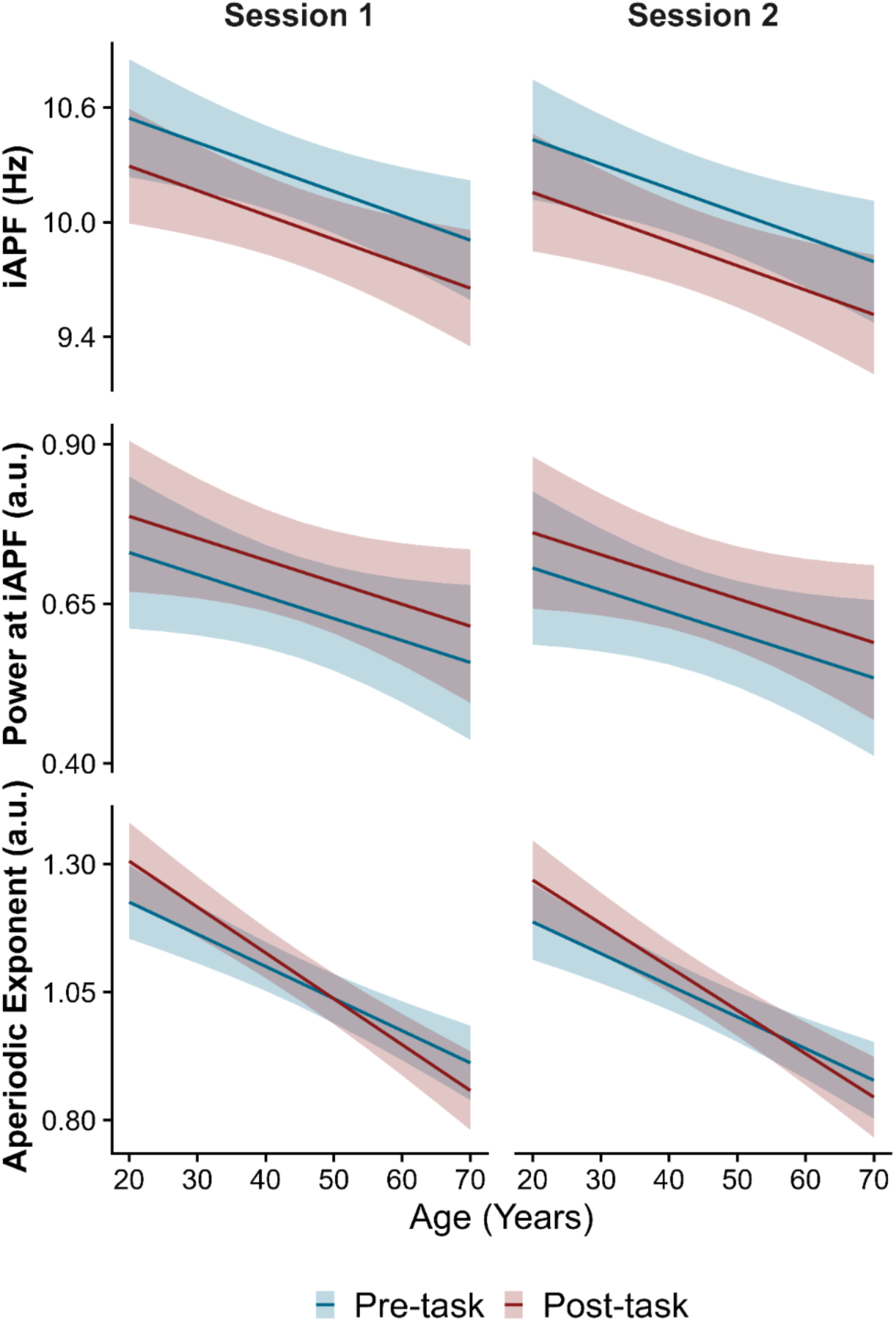
Predicted longitudinal lifespan trajectories of periodic and aperiodic rsEEG markers. Trajectories are based on the winning linear mixed-effects model for each metric (Table S4; Table 3), fitted to the combined longitudinal dataset (encompassing both Session 1 and Session 2) to represent the predicted marginal effects of chronological age (x-axis) on the electrophysiological markers (y-axis), whilst accounting for the effect of the five-year follow-up period. The continuous lines and color-matched shaded ribbons depict these predictions and their 95% confidence intervals, comparing pre-task (blue) and post-task (red) recordings. The left and right columns represent predictions for Session 1 and Session 2 data, respectively. Vertically, the panels depict trajectories for individual alpha peak frequency (iAPF; top), its associated power (middle), and the aperiodic exponent (bottom).

#### Power at iAPF

As for iAPF, Model L4 (*R^2^m* = 0.24, *R^2^c* = 0.88) best explained power at the iAPF (Table S5). We found no significant within-participant effect of session (*b* = -0.02, *SE* = 0.02, *t*(100) = -1.56, *p* = 0.12) nor a significant cross-sectional association with age (*b* = -0.003, *SE* = 0.002, *t*(100) = -1.87, *p* = 0.06), although the latter approached statistical significance. Similarly to previous results, power was significantly higher post-compared to pre-task (*b* = 0.06, *SE* = 0.01, *t*(100) = 4.39, *p* < 0.001; Fig. 6) as well as in the eyes-closed compared to the eyes-open conditions (*b* = 0.40, *SE* = 0.03, *t*(100) = 14.79, *p* < 0.001). There was no significant main effect related to sex (*b* = -0.05, *SE* = 0.05, *t*(100) = -0.83, *p* = 0.41).

#### Aperiodic exponent

For the aperiodic exponent, the best-fitting model (Model L2; *R^2^m* = 0.31, *R^2^c* = 0.87) included the same fixed effects as the above models as well as an additional interaction term between age and task (Table S5). We found that the exponent significantly decreased from session 1 to session 2 (*b* = -0.03, *SE* = 0.01, *t*(100) = -2.83, *p* = 0.006). The remaining pattern of results confirmed the outcomes from the session-wise models (Fig. 6). Specifically, we found that the exponent was significantly steeper in eyes-closed compared to eyes-open recordings (*b* = 0.11, *SE* = 0.01, *t*(100) = 11.50, *p* < 0.001) and there was no significant effect of sex (*b* = 0.006, *SE* = 0.03, *t*(100) = 0.19, *p* = 0.85). Importantly, the significant task-by-age interaction (*b* = -0.002, *SE* = 0.0006, *t*(100) = - 4.40, *p* < 0.001) was further evidenced, superseding the standalone main effects of age (*b* = -0.006, *SE* = 0.001, *t*(100) = -5.87, *p* < 0.001) and task (*b* = 0.13, *SE* = 0.03, *t*(182) = 4.90, *p* < 0.001).

As in the session-wise analyses, post-hoc tests revealed that even though both the pre- (*b* = -0.006, *SE* = 0.001, *t*(103) = -5.67, *p* < 0.001) and post-task (*b* = -0.009, *SE* = 0.001, *t*(103) = -7.33, *p* < 0.001) exponents were significantly and negatively related to age, the latter effect was significantly stronger (pre-post slope difference = 0.002, *SE* = 0.0006, *t*(102) = 4.27, *p* < 0.001). Furthermore, we investigated the effects of the task at each 10-year lifespan milestone between the ages of 25 and 65. In accordance with the Session 1 and 2 results, the exponent was significantly steeper post-compared to pre-task for younger cohorts at the ages of 25 (pre-post contrast estimate = -0.07, *SE* = 0.01, *t(*102) = -4.78, *p* < 0.001), 35 (pre-post contrast estimate = -0.04, *SE* = 0.01, *t*(102) = -4.26, *p* < 0.001), and 45 (pre-post contrast estimate = -0.02, *SE* = 0.008, *t*(102) = -2.21, *p* = 0.03). There was no difference in the exponent between pre- and post-task recordings at the ages of 55 (pre-post contrast estimate = 0.006, *SE* = 0.01, *t*(102) = 0.59, = 0.56), whilst a significantly decrease was observed at the age of 65 (pre-post contrast estimate = 0.03, *SE* = 0.01, *t*(102) = 2.14, *p* = 0.04).

### Validation on an independent, ageing cohort

To confirm that our findings generalise beyond the specific task architecture of the Dortmund dataset, we evaluated changes in the three EEG measures of interest in an independent cohort of healthy, older adults, who completed a different cognitive paradigm. Additionally, the narrow age range of this sample (63-77 years) provided an ideal framework to assess task-related dynamics independently of continuous age trends. For pre- and post-task grand average power spectra and scalp topographies of three examined rsEEG metrics, see Fig. S2. Table 6 shows the means are standard deviations related to each spectral metric separately for pre- and post-task recordings.

**Table 6.** Descriptive statistics for spectral metrics in the validation cohort of healthy older adults (*N* = 71). Within each cell, the mean (standard deviation) is shown for individual alpha peak frequency (iAPF), power at iAPF, and the aperiodic exponent, separately pre-task and post-task eyes-open recordings.

|  | iAPF<br>(Hz) | Power<br>at iAPF (a.u.) | Aperiodic<br>Exponent (a.u.) |
| --- | --- | --- | --- |
| Pre-task | 10.10<br>(1.00) | 0.61<br>(0.29) | 0.94<br>(0.18) |
| Post-task | 9.89<br>(0.98) | 0.671<br>(0.30) | 0.91<br>(0.19) |

We tested two different linear mixed-effects model variants on each EEG metrics of interest (Table S6). Both included fixed effects for task (pre- or post-task), age, sex as well as random intercepts for participants. The more complex model additionally included an interaction between age and time. For each frequency measure, the simpler model without the time-age interaction provided a better model fit based on the BIC (Table S6; iAPF: *R^2^m* = 0.07, *R^2^c* = 0.83; power at iAPF: *R^2^m* = 0.03, *R^2^c* = 0.92; aperiodic exponent: *R^2^m* = 0.02, *R^2^c* = 0.81). For an overview of the parameter estimates and related *p*-values from each winning model, see Table 7.

**Table 7.** Parameter estimates and associated *p*-values in the independent, validation cohort of healthy, older adults. Coefficients (*b*) and *p*-values (*p*) are presented for individual alpha peak frequency (iAPF), power at iAPF, and the aperiodic exponent. Data reflect eyes-open resting-state recordings only. Optimal model structures were selected based on the Bayesian Information Criterion (BIC), with full model comparison details provided in Table S6. Reference levels for categorical predictors are pre-task for task (post-task vs. pre-task), closed or eye condition (closed vs. open), and male for sex (female vs. male). Age was a continuous predictor between 63 and 77 years. We observed significant task-induced changes in all three rsEEG metrics.

|  | iAPF |  | Power at iAPF |  | Aperiodic Exponent |  |
| --- | --- | --- | --- | --- | --- | --- |
|  | b | p | b | p | b | p |
| Intercept | 6.38 | 0.011 | 0.31 | 0.697 | 1.42 | 0.003 |
| Task | -0.21 | 0.003 | 0.10 | <0.001 | -0.03 | 0.017 |
| Age | 0.05 | 0.148 | 0.004 | 0.695 | -0.01 | 0.323 |
| Sex | 0.48 | 0.058 | 0.48 | 0.058 | -0.02 | 0.702 |

Our analyses indicated significant task-related changes in all metrics (Fig. S3). Specifically, iAPF (*b* = 0.-21, *SE* = 0.07, *t*(70) = -3.00, *p* = 0.004) and the aperiodic exponent (*b* = -0.03, *SE* = 0.01, *t*(70) = -2.41, *p* = 0.02) decreased significantly from pre- to post-task, whilst there was a significant task-induced increase in alpha power (*b* = 0.10, *SE* = 0.01, *t*(70) = 7.05, *p* < 0.001). However, we found no significant effect of age (iAPF: *b* = 0.05, *SE* = 0.04, *t*(68) = 1.46, *p* = 0.15, alpha power: *b* = 0.004, *SE* = 0.01, *t*(68) = 0.39, *p* = 0.70, aperiodic exponent: *b* = -0.07, *SE* = 0.07, *t*(68) = -0.99, *p* = 0.32) or sex (iAPF: *b* = 0.48, *SE* = 0.26, *t*(68) = 1.91, *p* = 0.06, alpha power: *b* = -0.02, *SE* = 0.08, *t*(68) = -0.23, *p* = 0.82, aperiodic exponent: *b* = -0.02, *SE* = 0.05, *t*(68) = -0.38, *p* = 0.70).

## Discussion

By leveraging large-scale cross-sectional, longitudinal, and independent validation cohorts, our study demonstrates that acute cognitive exertion systematically modulates periodic and aperiodic rsEEG markers of ageing and pathology, including the iAPF, its associated power, and the aperiodic exponent. Cross-sectionally, all examined electrophysiological metrics exhibited significant age-related decreases across the adult lifespan, a pattern corroborated longitudinally by significant five-year within-participant reductions in both iAPF and the aperiodic exponent. Importantly, we observed a consistent interaction between age and task-induced shifts in the aperiodic exponent, characterised by a significantly stronger effect between the aperiodic exponent and chronological age post-task compared to pre-task. This suggests that electrophysiological states captured following a cognitive challenge may provide a more sensitive reflection of neurophysiological ageing than traditional, pre-task recordings.

Cross-sectionally, all examined rsEEG metrics exhibited significant age-related changes across the adult lifespan. Consistent with previous studies^11,13,18,20,22,61^ both iAPF and the aperiodic exponent demonstrated a negative relationship with age in the cross-sectional session analyses as well as within participants over the 5-year interval. In alignment with reported non-linear lifespan dynamics^10,11,13,14^, iAPF additionally displayed a quadratic trajectory in Session 1, whereby it increased at age 25 followed by an accelerated decline from age 45 onwards. However, this quadratic effect was absent in Session 2 and the reduced Session 1 sub-sample, likely due to sample size constraints limiting power to detect nonlinear trends. Evidence for age-related declines in iAPF power was inconsistent; whilst significant in Session 1 and marginal in Session 2 (*p* = .04), the longitudinal model showed no significant cross-sectional effect (*p* = .06) or within-person change over 5 years. This pattern mirrors mixed findings between age and relative alpha power^11,18,51^, which may stem from subtle effect sizes that remain difficult to detect in moderately powered samples.

We also observed consistent, significant effects of eye condition across all rsEEG metrics. Consistent with previous studies, we found lower iAPF^7,33^ and higher associated power^7,10,33,61^ in eyes-closed recordings. This pattern reflects the established inverse relationship between alpha amplitude and peak frequency^62^, which has been linked to sensory processing and cortical inhibition (for a review, see Mierau et al.^62^). Notably, task-induced alpha power changes occurred exclusively during eyes-open recordings, suggesting that eyes-closed alpha may be less susceptible to transient state shifts, which stability is corroborated by the improved test-retest reliability of alpha power under eyes-closed conditions^6,63^. Similarly to previous research^7,61^, we observed steeper aperiodic exponents during eyes-open conditions, supporting the view that background aperiodic activity tracks contextual changes^61^ and may reflect E/I balance shifts^21,22^. Altogether, these findings emphasise the importance of assessing rsEEG metrics across both eye conditions^4^ and suggest that eyes-open recordings may show an increased sensitivity to task-induced state shifts.

In the periodic domain, we found a consistent task-induced and lifespan-wide decrease in iAPF in both eye conditions, alongside an associated power increase in eyes-open recordings. Previous studies examining task-related changes in alpha metrics are sparse and provided mixed results. iAPF has been reported to show task-related decreases^28,31^, increases^34^, as well as null results^33^ in young cohorts. The task-induced rise in power at iAPF aligns with previous reports of absolute^28^ and relative^30^ alpha power increases in young adults. In older cohorts, existent evidence indicates task-related reductions in absolute alpha power^32^ and an increase in relative alpha power^30^. By isolating true periodic alpha oscillations from the underlying aperiodic background, evaluating eyes-open and eyes-closed recordings, and demonstrating generalisability across different paradigms, our study clarifies these mixed findings. Ultimately, our results demonstrate that post-task alpha dynamics, characterised by persistent frequency slowing alongside power increase in the eyes-open recordings, represent a general lifespan-wide neurophysiological phenomenon.

In the aperiodic domain, we observed a divergent lifespan trajectory, whereby acute cognitive exertion induced a post-task steepening of the aperiodic exponent in young adults, which progressively flattened with increasing age until a net decrease was reached at the age of 65. This effect was successfully replicated in the independently acquired validation cohort of older adults (63-77 years). Importantly, these task-induced changes resulted in a stronger negative age-related effect (i.e., steeper marginal slopes) post-task compared to pre-task in both the session-wise and longitudinal analyses. To our knowledge, Yang et al.^30^ represents the sole study evaluating task-related shifts in the exponent, which analyses showed a similar interaction between task and age groups. By modeling age continuously across the lifespan and validating our findings across different eye conditions and cohorts, we demonstrate that these age-dependent shifts across the lifespan are robust across experimental conditions.

The dissociation between task-induced alpha shifts and age-dependent exponent trajectories indicates that these spectral features index distinct physiological mechanisms across the lifespan^10,11^. The uniform task-induced decreases in iAPF and concurrent increases in power may signal increased mental fatigue^64,65^ or top-down inhibition^66,67^. At the same time, task-related shifts in the exponent could reflect age-related changes in cortical E/I balance^21,22,68^, vigilance^69^, or synaptic kinetics^68^.

Whilst the task-induced volatility in these electrophysiological markers poses a challenge for biomarker research requiring stable, trait-like measures, the consistency of these shifts across cohorts and paradigms highlights their systematic nature. These findings raise the important methodological question of whether pre- or post-task resting state serves as a better proxy for trait-level neural features. Our results demonstrate a stronger age effect in the post-task compared to the pre-task aperiodic exponents, parallelling a growing body of rsEEG studies indicating that post-task metrics exhibit superior clinical discrimination ability compared to traditional pre-task baselines^32,34,36,37^.

Post-task recordings may uniquely function as a ‘stress test’, analogous to monitoring cardiovascular dynamics after physical exertion^34,35^, whereby intrinsic traits are more clearly expressed during recovery. Underlying mechanisms may include the neural carry-over of task-related activation^70^ potentially reflecting compensatory activation^36^, varying individual delays in the homeostatic return to baseline resting-state^36,37,71^, or post-task performance self-appraisal^36,71^. Alternatively, the post-task advantage may stem from the standardisation of participants’ preceding experiential and neurophysiological states. Pre-task recordings may reflect a random distribution of pre-experimental states, such as varying levels of arousal, emotional tone, or caffeine intake prior to entering the laboratory, which may introduce noise that can obscure individual trait differences. By introducing a uniform experience, cognitive exertion could minimise state variance and thus allow trait-level characteristics to emerge more reliably. Ultimately, elucidating the precise neurophysiological mechanisms behind these task-induced shifts will be essential for validating the post-task framework in biomarker research.

Several limitations of this study should be noted. First, whilst computing a global electrode averages ensures cross-dataset reproducibility and provides a topographically unbiased representation of both posterior-dominant alpha and central-parietal aperiodic metrics, this may also obscure localised dynamics. High-density investigations evaluating spatially specific modulations will be critical to capturing localised patterns of age-related changes. Second, transient state variables (e.g., sleep quality, caffeine intake, mood, or arousal) were not systematically measured, leaving their precise impact on these spectral metrics unaddressed. To build on these findings, future work could track post-task recovery trajectories to determine whether the duration required to return to pre-task baselines varies systematically across the lifespan. Additionally, studies could also dissociate whether active cognitive exertion and passive state harmonisation (e.g., watching a neutral film) produce comparable shifts in spectral metrics. Evaluating whether post-task metrics offer superior discrimination across clinical and cognitive traits will also be essential for establishing the utility of the post-task framework for translational neuroscience. Finally, building on these studies and utilising large-scale data sets, explainable machine-learning models could be developed for data-driven characterisation, classification, and prediction of healthy and pathological conditions.

In summary, our study demonstrates that acute cognitive exertion systematically modulates periodic and aperiodic resting-state EEG markers of healthy and pathological ageing across the adult lifespan. As evidenced by the divergent task-induced trajectories of the aperiodic exponent across the lifespan, our results suggest that post-task recordings offer a more sensitive measure of neurophysiological ageing than traditional pre-task recordings. These findings support the application of post-task paradigms as a methodological framework to evaluate neurophysiological dynamics in lifespan, individual differences, and clinical research.

## Data availability

Supporting data are available from the corresponding authors upon reasonable request.

## Code availability

The resting-state data pre-processing and spectral decomposition scripts are openly available on GitHub (https://github.com/RCNS-BIC/rsEEG_BANetal2026).

## Acknowledgements

This work was supported by the National Brain Research Program 3.0. by the Hungarian Academy of Sciences (NAP2022-I-1/2022; PI: Z.V.); the Hungarian Research Network (HUN-REN; 298/4/2023/HF; PI: Z.V.), and the National Research, Development and Innovation Fund (No. 2024-1.1.2-NAGYVÁLL_FÓKUSZ-2025-00001; PI: Z.V.). We express our sincere gratitude to Anna Kocsis, István Hevesi, and Vivien Mária Lesku for their invaluable assistance with the data collection of the validation dataset.

## Author contributions

K.B.: conceptualisation, data curation, project administration, formal analysis, investigation, methodology, visualisation, writing – original draft, writing – review & editing. B.W.: conceptualisation, formal analysis, methodology, software, supervision, writing – review & editing. T.A.: data curation, software, writing – review & editing. P.D.G.: data collection, data curation, investigation, writing – review & editing. E.W.: data curation, investigation. Z.V.: conceptualisation, funding acquisition, supervision, project administration, writing – review & editing.

