## Supplementary information for "Cognitive exertion reshapes resting-state EEG markers across the adult lifespan"

#### Eye-tracking data acquisition for the validation dataset

The Eyelink 1000 Plus (SR Research Ltd., Canada) eye-tracker recorded participants' binocular eye movements and pupil area at a sampling rate of 1000 Hz. The eye-tracking camera was positioned at the base of the monitor, securing an unobstructed view of the screen. Prior to recording, gaze position was mapped to screen coordinates using the standard EyeLink 9-point grid calibration procedure. Calibration was repeated where visual inspection of the fixation targets indicated unstable or inaccurate gaze tracking.

### Supplementary tables

**Table S1.** Information on the number of participant-wise rejected independent components, clean epochs, excluded channels, and group FOOOF model fit in the different Dortmund Vital Study datasets. Within each cell, data are mean (*SD*). Further details on the independent component rejection criteria can be found in Methods.

| Dataset | | | Independent components Rejected | Epochs after pre-processing | FOOOF model fit | Excluded channels |
|---|---|---|---|---|---|---|
| Session 1 | Eyes-Open | Pre-task | 33.82 (7.21) | 60.15 (21.04) | 0.97 (0.03) | 2.88 (7.86) |
| | | Post-task | 31.06 (7.22) | 52.79 (21.78) | 0.97 (0.03) | 3.04 (8.21) |
| | Eyes-Closed | Pre-task | 26.25 (8.17) | 80.13 (13.37) | 0.98 (0.01) | .60 (2.55) |
| | | Post-task | 22.82 (7.60) | 75.34 (16.17) | 0.98 (0.01) | .43 (2.16) |
| Session 2 | Eyes-Open | Pre-task | 34.97 (7.63) | 60.01 (22.01) | 0.96 (0.03) | 3.70 (9.33) |
| | | Post-task | 31.06 (8.07) | 52.20 (23.40) | 0.96 (0.04) | 4.23 (10.73) |
| | Eyes-Closed | Pre-task | 26.12 (7.45) | 79.82 (12.14) | 0.98 (0.01) | 0.29 (1.12) |
| | | Post-task | 23.22 (7.72) | 747.34 (14.56) | 0.98 (0.01) | 0.14 (.45) |

**Table S2.** Information on the number of participant-wise rejected independent components, clean epochs, excluded channels, and group FOOOF model fit in the validation dataset of healthy, older adults.. Within each cell, data are mean (*SD*). Further details on the independent component rejection criteria can be found in Methods.

| Data | Independent components Rejected | Epochs after pre-processing | FOOOF model fit | Excluded channels |
|---|---|---|---|---|
| Pre-task | 29.01 (6.68) | 70.62 (21.16) | 0.96 (0.03) | 4.11 (9.83) |
| Post-task | 28.95 (6.22) | 66.90 (21.49) | 0.96 (0.03) | 3.30 (6.71) |

**Table S3. Model comparison in the Session 1 dataset.** Bayesian Information criterion (BIC) for each model variant predicting each EEG metric of interest. Lower values illustrate a better model fit. The winning model is marked bold.

| Model | iAPF | Power at iAPF | Aperiodic Exponent |
|---|---|---|---|
| Model 1:<br>~ eye * task * age + sex +<br>(1 \| participant) | 3385.86 | 474.26 | -1503.39 |
| Model 2:<br>~ eye * task * poly(age,2) + sex + (1 \|<br>participant) | 3401.43 | 494.40 | -1480.77 |
| Model 3:<br>~ eye * task * age + sex +<br>(eye + task\| participant)) | 3288.69 | 198.84 | **-1569.24** |
| Model 4:<br>~ eye * task * poly(age,2) + sex + (eye<br>+ task \| participant) | 3301.49 | 221.38 | -1548.28 |
| Model 5:<br>~ eye * task + age + sex +<br>(1 \| participant) | 3375.59 | 458.57 | -1470.06 |
| Model 6:<br>~ eye * task + poly(age,2) + sex + (1\|<br>participant) | 3372.16 | 465.80 | -1463.84 |
| Model 7:<br>~ eye * task + age + sex +<br>(eye + task \| participant) | 3278.72 | **182.77** | -1542.89 |
| Model 8:<br>~ eye * task + poly(age,2) + sex +<br>factor(sex) + (eye + task \| participant) | **3273.96** | 188.88 | -1538.17 |

**Table S4. Longitudinal reliability of spectral measures.** Intra-class correlation (ICC) coefficients measuring absolute agreement (A-1) were computed between Session 1 and Session 2 (5 years apart) for the stratified, longitudinal sub-sample (N = 100) of the Dortmund Vital Study. ICCs were quantified for individual alpha peak frequency (iAPF), its associated power (power at IAPF), and the aperiodic exponent. Within each cell, data are ICC coefficients (*r*).

| | | iAPF | Power at iAPF | Aperiodic exponent |
|---|---|---|---|---|
| Eyes- Open | Pre-task | 0.62 | 0.75 | 0.78 |
| | Post-task | 0.63 | 0.76 | 0.75 |
| Eyes- Closed | Pre-task | 0.83 | 0.91 | 0.76 |
| | Post-task | 0.84 | 0.73 | 0.78 |

**Table S5. Model comparison in longitudinal dataset containing data from Session 1 and 2 of the Dortmund Vital Study.** Bayesian Information criterion (BIC) for each model variant predicting each EEG metric of interest. Lower values illustrate a better model fit. The winning model is marked bold.

| Model | iAPF | Power at iAPF | Aperiodic Exponent |
|---|---|---|---|
| Model 1:<br>~ session * task * age + sex + eye + (eye + session + task\| participant) | 1591.69 | -9.41 | -922.57 |
| Model 2:<br>~ session + task * age + sex + eye + (eye + session + task \| participant) | 1571.96 | -24.61 | **-941.34** |
| Model 3:<br>~ session * task + age + sex + eye + (eye + session + task \| participant) | 1572.36 | -23.56 | -924.54 |
| Model 4:<br>~ session + task + age + sex + eye + (eye + session + task \| participant) | **1565.85** | **-30.24** | -930.33 |

**Table S6. Model comparison the independently acquired, elderly dataset.** Bayesian Information criterion (BIC) for each model variant predicting each EEG metric of interest. Lower values illustrate a better model fit. The winning model is marked bold.

| Model | iAPF | Power at iAPF | Aperiodic Exponent |
|---|---|---|---|
| Model 1:<br>~ task * age + sex + (1 \| participant) | 344.69 | -42.27 | -114.88 |
| Model 2:<br>~ task + age + sex + (1 \| participant) | **342.66** | **-43.24** | **-129.50** |

### Supplementary figures

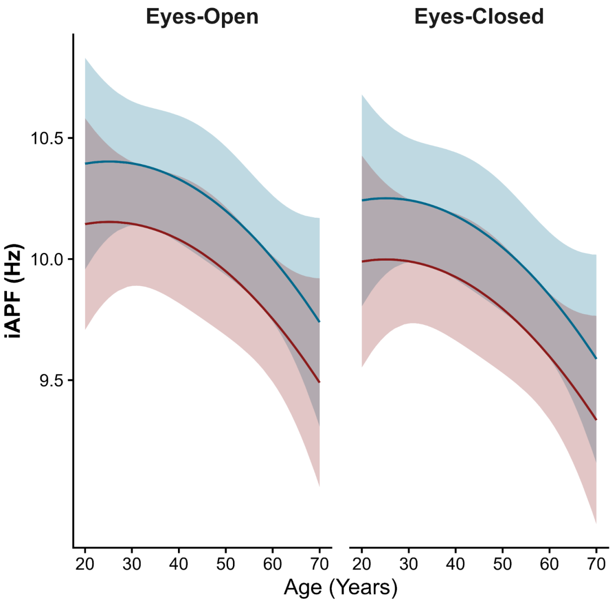


**Figure S1. Predicted lifespan trajectories of iAPF in Session 1, restricted to the Session 2 sub-sample.** We fitted the winning linear mixed-effects model, established in Session 1, to the sub-sample of Session 1 data (N = 100) with matching participants from Session 2. Trajectories represent the predicted marginal effects of chronological age (x-axis) on iAPF (y-axis). The continuous lines and color-matched shaded ribbons depict these predictions and their 95% confidence intervals, comparing pre-task (blue) and post-task (red) recordings. The left and right columns represent predictions for eyes-open and eyes-closed recordings, respectively.

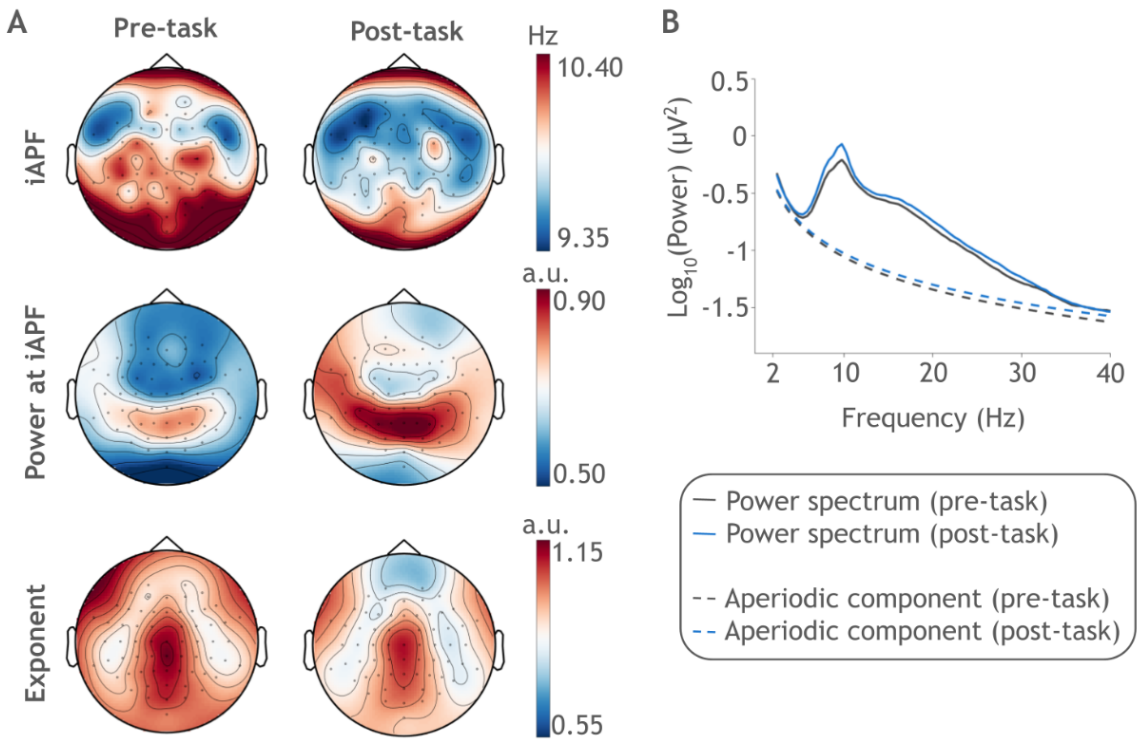


**Figure S2. Scalp topographies and power spectra before and after the cognitive task battery for the validation, ageing dataset. A,** The left and right side shows pre- and post-task topographies, respectively, averaged across participants. Horizontally, the topographies depict individual alpha peak frequency (iAPF; top), its associated power (middle), and the aperiodic exponent (bottom). **B,** Mean raw power spectra (continuous lines) and aperiodic components (dashed lines) compared between pre-task (grey) and post-task (blue) recordings, averaged across all channels and participants.
